# SSNA1 organizes the distal luminal centriolar network and promotes ciliogenesis without microtubule association

**DOI:** 10.1101/2025.04.28.648957

**Authors:** Yi-Chi Huang, Xiao-Jing Chong, Ting-Jui Ben Chang, Emma van Grinsven, Wei-Ju Chen, Wen-Bin Hsu, Jeffrey M. Beekman, Tang K. Tang, Anna Akhmanova, T. Tony Yang, Jen-Hsuan Wei

## Abstract

The distal lumen of the centriole plays a critical role in ciliogenesis, yet the molecular composition, spatial organization, and targeting hierarchy of distal luminal proteins remain poorly understood. In this study, we identify Sjögren’s syndrome nuclear autoantigen 1 (SSNA1) as a *bona fide* centriolar protein and a key regulator of ciliogenesis in mammalian cells. Using our newly developed knockout (KO)-validated antibody, we show that, contrary to previous reports, SSNA1 does not reside in the nucleus, midbody, or ciliary axoneme. Instead, super-resolution imaging combined with expansion microscopy (ExM) reveals that SSNA1 localizes to the distal lumen of centrioles and the basal bodies of both primary and motile cilia, where it is arranged in a ring-like configuration with 9-fold symmetry and apart from centriolar microtubules. Molecular dissection using tag-free SSNA1, its oligomerization-deficient mutants, and microtubule co-pelleting assays further demonstrates that SSNA1, previously described as a microtubule nucleator, stabilizer, and branching factor, does not bind microtubules *in vitro*. Interactor screening and KO analysis unveil a hierarchical targeting network involving a C2CD3-SSNA1-LRRCC1 axis in the distal lumen. Functional characterization indicates that although dispensable for cell division, overall centriole organization and duplication, SSNA1 promotes cilia assembly by facilitating CP110 removal. Our findings redefine the physiological role of SSNA1 as part of the distal luminal module contributing to ciliogenesis and provide new insights into the molecular architecture and functional relevance of the distal centriolar lumen.

**Significance:** Centrioles template the assembly of cilia and flagella, whose dysfunction causes human diseases known as ciliopathies. Key centriolar structures - the cartwheel in the proximal lumen, the inner scaffold in the central lumen, and the distal and subdistal appendages outside the distal lumen - have been extensively characterized. However, the molecular organization and functional relevance of the distal lumen remain largely unexplored. Here, we identify SSNA1 as a novel centriolar component at the distal lumen and an essential regulator of ciliogenesis. Our work challenges the long-held view of SSNA1 as a microtubule-associated protein and provides new insights into the architecture, interaction network, targeting hierarchy, and function of the distal centriolar lumen, exemplifying how it modulates centriolar organization and drives ciliogenesis.

## Introduction

Centrosomes are evolutionarily conserved microtubule-based organelles that serve as templates for the assembly of cilia and flagella (1). Each centrosome consists of two centrioles surrounded by pericentriolar material (PCM). In mammalian cells, centrioles are built by microtubule triplets (2, 3), which are arranged in a cylindrical fashion with 9-fold radial symmetry, forming a barrel-shaped structure with a central lumen. Microtubule triplets extend from the proximal base of the centriole to its distal end, where they are converted into microtubule doublets that elongate further to form the ciliary axoneme during ciliogenesis (4, 5).

Centrioles possess an inherent polarity along their proximal-distal axis. Based on their structural composition and functional roles, centrioles can be divided into three distinct regions: proximal, central, and distal (6–8). The proximal end is involved in centriole cohesion (9) and recruitment of the PCM (10–13), which nucleates microtubules for spindle assembly during cell division (14). In addition, within the proximal lumen lies the cartwheel (15–17), a key structural element essential for the 9-fold radial symmetry (18–20) and duplication of centrioles (21–23). The central domain, or inner scaffold (24–27), occupies most of the luminal volume and is critical for maintaining centriole length and structural integrity. Finally, the distal region is best known for its role in assembling cilia and flagella. This function has been largely attributed to two specialized structures, known as the distal appendage (DA) and the subdistal appendage (SDA) (28–30), which are attached to the outer wall of the mother centriole. While DAs enable centriole docking to preciliary vesicles (PCVs) or the apical plasma membrane for axoneme initiation (31), SDAs anchor microtubules and regulate cilia positioning (32).

In addition to axial polarity, centriolar proteins also exhibit differential radial distributions, forming a highly ordered and concentric macromolecular assembly. The exquisite radial organization is best exemplified by the hierarchical arrangement of DA and SDA proteins on the mother centriole (29). Recent breakthroughs in CRISPR-mediated gene editing and advanced imaging techniques including super-resolution microscopy and ExM have illuminated how DA and SDA proteins are sequentially recruited and assembled at the outer wall of centrioles (29, 33–36), shedding light on their intricate architecture and the stepwise pathways that guide their centriolar targeting.

Despite these recent findings, the distal lumen remains an enigmatic part of the centriole (37). Given its close proximity to the sites where distal appendage vesicles (DAVs) fuse to form the ciliary vesicle (CV) (38, 39) and where the axoneme begins to grow, the distal lumen may also play a role in ciliogenesis. To date, only a limited number of distal luminal proteins have been identified, including C2CD3 (8, 40, 41), LRRCC1 (42, 43), and SFI1 (8, 44). Indeed, depletion of these proteins impair cilia assembly (40–44). However, due to the lack of a molecular inventory, an undefined targeting hierarchy, and visualization challenges imposed by the diffraction limit of conventional light microscopy, a comprehensive view of the components, organization, and assembly pathway at the distal lumen remains lacking, leaving a substantial gap in our understanding of its molecular architecture and functional relevance. Therefore, identifying novel distal luminal proteins, analyzing their spatial organization, and deciphering their association network represents a crucial first step towards unraveling the molecular mechanisms of ciliogenesis.

Here, we report Sjögren’s syndrome nuclear autoantigen 1 (SSNA1) as a centriolar protein that specifically localizes to the distal lumen. SSNA1, also known as nuclear antigen of 14 kDa (NA14), is a small coiled-coil protein originally identified as a target of autoantibodies from a patient with Sjögren’s syndrome (45). Although cloned from a human testis cDNA library, SSNA1 is expressed in multiple tissues (45). Evolutionarily, it is highly conserved across multicellular organisms, ranging from nematodes, fish, frogs and mice to humans, as well as occurring in unicellular organisms, including *Chlamydomonas* (46, 47), trypanosomes (48) and toxoplasmas (49). Despite its name, SSNA1 has also been reported to localize to several microtubule-rich structures outside the nucleus, including centrosomes and midbodies, where it was proposed to regulate cell division and cytokinesis (50, 51). In *Chlamydomonas*, its homolog DIP13 was found at basal bodies and appeared to partially decorate the flagellar axoneme (46).

Several *in vitro* studies showed that recombinant SSNA1 has the ability to oligomerize and self-assemble into filaments (48, 49, 52–54). Recently, SSNA1 has been further characterized as a microtubule-associated protein (MAP) that binds longitudinally along microtubules and promotes microtubule branching *in vitro* (53, 54). When overexpressed in cultured neurons, SSNA1 was found to accumulate at axonal branching sites (53) where it promotes axon extension and branching (51, 53) through SSNA1-induced microtubule branching. In contrast, another study using single-molecule total internal reflection fluorescence (TIRF) microscopy did not detect microtubule branching and suggested that SSNA1 modulates microtubule dynamics by slowing its growth, shrinkage, and catastrophe while promoting rescue (55). Notably, despite the reported *in vitro* binding of SSNA1 to microtubules, direct association of SSNA1 with microtubules has not been observed *in vivo*.

Given the multiple reported localizations of overexpressed SSNA1, we undertook a systematic investigation to determine its subcellular localization at the endogenous level and to analyze its physiological function in mammalian cells. Using a newly developed KO-validated antibody, we demonstrate that SSNA1 does not reside in the nucleus, at the midbody, or along the ciliary axoneme as previously reported. Instead, super-resolution imaging combined with ExM reveals that SSNA1 localizes specifically to the distal lumen of centrioles and is present at the basal bodies of both primary and motile cilia. In the distal lumen, SSNA1 is arranged in a 9-fold ring configuration and distant from centriolar microtubules. Using tag-free recombinant SSNA1, its newly generated oligomerization-deficient mutants, and microtubule co-pelleting assays, we show that previous results identifying SSNA1 as a MAP were due to non-specific binding of the His tag to microtubules *in vitro*. Through interactor screening and KO analysis in cells, we further unveil a hierarchical targeting and interaction network involving a C2CD3-SSNA1-LRRCC1 axis in the distal lumen. Functional analysis indicates that SSNA1 is dispensable for appendage assembly, centriole duplication and cell division but promotes cilia assembly by facilitating the removal of the ciliogenesis suppressor CP110 from the mother centriole. In conclusion, our findings redefine the role of SSNA1, resolve discrepancies, and uncover its cellular function in organizing the distal centriolar lumen and promoting ciliogenesis.

## Results

### Generation of SSNA1 KO cell lines and KO-validated SSNA1-specific monoclonal antibody

SSNA1 has been reported to localize to several distinct subcellular compartments, including the nucleus (45) and various microtubule-rich structures, such as centrosomes and midbodies in dividing mammalian cells (46, 51), basal bodies and flagellar axonemes in green algae (46), and axonal branching sites in neurons (53). However, some of the localizations appear to be inconsistent between species and studies, most of which have employed overexpression to analyze SSNA1 localization and function (45, 47, 51, 53, 56). We hypothesized that the discrepancies may be due to overexpression and therefore set out to define the subcellular localization and physiological function of endogenous SSNA1 in mammalian cells.

To this end, we first generated SSNA1 KO lines in human hTERT-immortalized retinal pigment epithelial-1 (RPE-1) cells using CRISPR/Cas9 genome editing (Fig. S1). We implemented two different KO designs using a double-cut strategy (Fig. S1*A*), which gave rise to two independent clones, KO#1 and KO#2. Both KO clones were validated by three complementary approaches: genomic DNA sequencing (Fig. S1*B*, assembled sequences are indicated by red bars at the top), Western blotting (Fig. S1*D*), and immunofluorescence (Fig. S2). Sanger sequencing results (Fig. S1*B*) and genotyping PCR (Fig. S1*C*) confirmed successful deletion of the targeted genomic regions, with the first cut placed upstream of the ATG start codon, resulting in no protein translation. Western blotting analysis of total cell lysates using commercially available rabbit polyclonal antibodies PTG further confirmed the absence of a 14 kDa band corresponding to SSNA1 in the KO cells (Fig. S1*D*). However, immunofluorescence staining of both SSNA1 KO clones with the same PTG antibodies still showed positive signals at centrosomes similar to those in wild-type (WT) cells (Fig. S2, only KO#2 is shown), indicating that the PTG antibodies cross-react with other centrosomal proteins (51).

To obtain SSNA1-specific antibodies for cellular and functional analyses, we generated antibodies against purified tag-free full-length human SSNA1. Despite several attempts to raise polyclonal antibodies in rabbits (Rb) and guinea pigs (GP), all of our efforts were unsuccessful (Fig. S2). All of these polyclonal antibodies cross-reacted with centrosomes by immunofluorescence in SSNA1 KO cells, similar to the commercial polyclonal antibodies PTG. Therefore, we turned to hybridoma technology to develop a mouse monoclonal antibody, which ultimately resulted in clone A6. This monoclonal antibody specifically recognized SSNA1, as the corresponding immunofluorescence signal completely disappeared in SSNA1 KO cells (Fig. S2). While all further localization and functional analyses were performed with both SSNA1 KO clones and showed consistent results, only SSNA1 KO#2 is shown as a representative unless otherwise indicated.

### SSNA1 is a bona fide centriolar protein

Using the newly developed and KO-validated monoclonal antibody A6, we examined the subcellular localization of SSNA1 in proliferating and resting RPE-1 cells (Fig. 1*A*). Exponentially growing and starvation-induced ciliated RPE-1 cells were stained with antibodies against SSNA1 (A6 and PTG, Figs. 1*A* and S3*A*, respectively), the centrosome marker CEP164, and acetylated tubulin (Ac-tub) that labels stable microtubules in interphase, spindle microtubules in mitosis, the midbody in cytokinesis, and the axoneme in ciliated cells. We observed that SSNA1 colocalized with CEP164 in both proliferating and resting cells, supporting that SSNA1 is a *bona fide* centrosomal protein. The centrosomal localization of SSNA1 persisted throughout the cell cycle (interphase, mitosis, and cytokinesis) as well as in the G0 phase of ciliated cells. However, the A6 monoclonal antibody did not detect SSNA1 at the midbody during cytokinesis (Fig. 1*A*, cytokinesis, circle), in contrast to PTG antibodies (Fig. S3*A*, circle), suggesting that the previously reported midbody localization (51) was caused by antibody cross-reactivity. Furthermore, we did not detect SSNA1 at the ciliary axoneme in RPE-1 cells (Fig. 1*A*, ciliated, circle), unlike in green algae where the SSNA1 homolog DIP13 appears to partially decorate the flagellar axoneme (46).

**Fig. 1:**
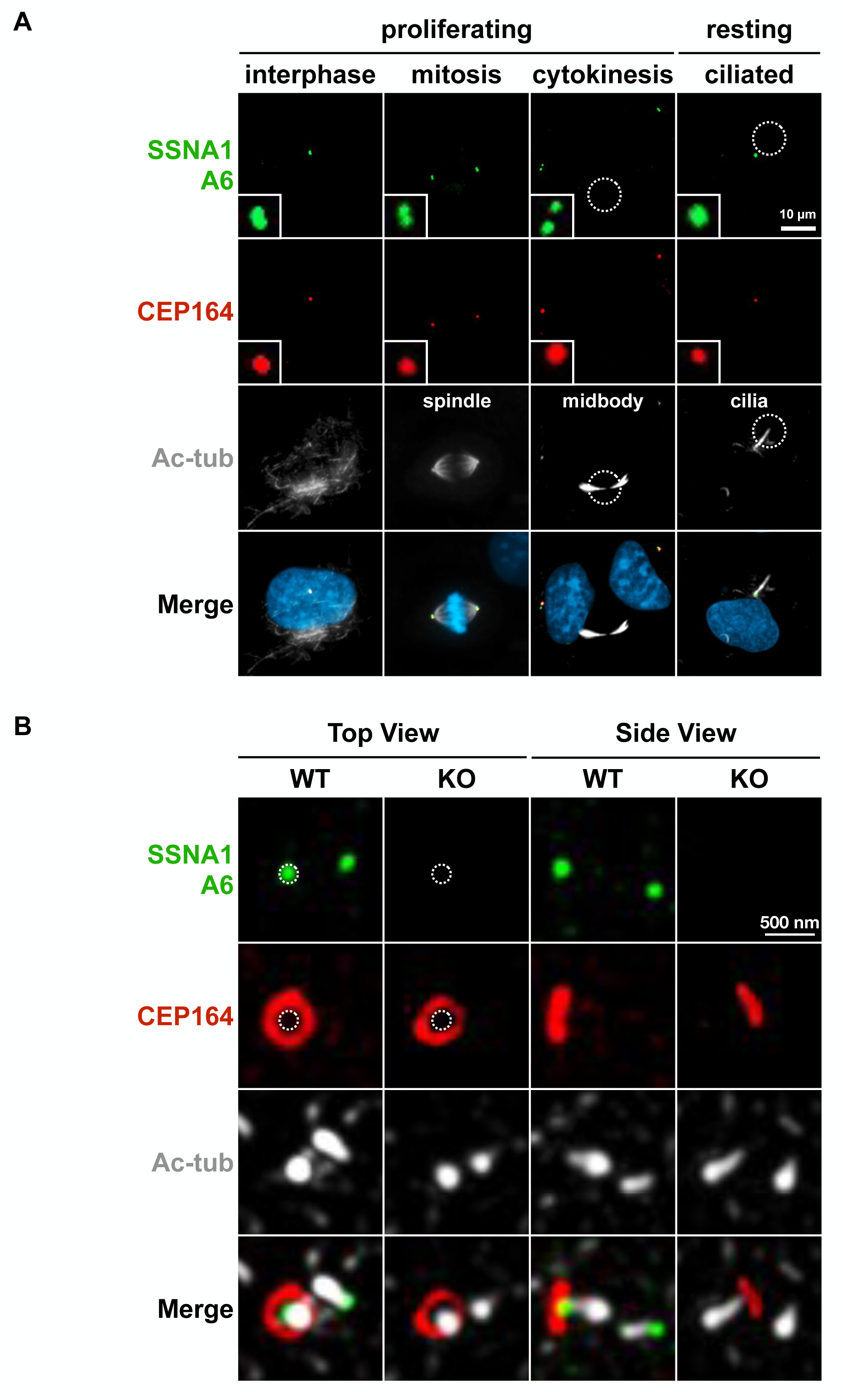
SSNA1 is a *bona fide* centriolar protein. **A,** Localization of SSNA1 in proliferating and resting RPE-1 cells. Exponentially growing and starvation-induced ciliated RPE-1 cells were stained with antibodies against SSNA1 (A6), the centrosomal marker CEP164, and acetylated tubulin (Ac-tub), which labels stable microtubules in interphase, spindle microtubules in mitosis, the midbody in cytokinesis, and the axoneme in ciliated cells. SSNA1 was found at centrosomes throughout the cell cycle and the basal bodies of primary cilia. 5x magnified images of the centrosomal region are shown in the insets. SSNA1 was not detected at the midbody (circled) during cytokinesis or at the ciliary axoneme (circled). Bar, 10 µm. **B,** Representative 3D-SIM images of the suborganellar localization of SSNA1 at centrosomes. Exponentially growing RPE-1 cells were stained with antibodies against SSNA1 (A6), the mother centriole/DA marker CEP164, and the centriolar microtubule marker acetylated tubulin (Ac-tub). A6 antibody identified two distinct puncta in WT cells, both of which colocalized with acetylated tubulin and were completely absent in KO cells. Top View: In WT cells, one SSNA1 signal was positioned within the CEP164 ring (circled) while the other was located at the end of the centriole, suggesting that SSNA1 localizes to the distal end of both mother and daughter centrioles. Side View: SSNA1 localized to the distal end with CEP164, confirming SSNA1 as a distal centriolar protein. Bar, 500 nm.

To investigate the suborganellar localization of SSNA1 at centrosomes, we used three-dimensional structured illumination microscopy (3D-SIM). RPE-1 WT and SSNA1 KO cells were stained with SSNA1 antibodies (A6 and PTG, Figs. 1*B* and S3*B*, respectively), together with markers for the mother centriole/DA (CEP164) and centriolar microtubules (acetylated tubulin, Ac-tub). In contrast to the epi-fluorescence results, 3D-SIM revealed multiple signals at centrosomes in WT cells labeled with PTG antibodies, with one spot located within the CEP164 ring and others appearing as satellite-like puncta around the two centrioles (Fig. S3*B*). In SSNA1 KO cells, the signal within the CEP164 ring was absent, indicating that this spot represents the true localization of SSNA1, while the surrounding puncta remained, indicating antibody cross-reactivity. In contrast, the A6 antibody labeled two distinct puncta in WT cells, both of which colocalized with the acetylated tubulin signal marking the centrioles. These two puncta disappeared in KO cells, further validating the specificity of the A6 antibody.

Notably, a top view of WT cells revealed that one SSNA1 signal was positioned within the CEP164 ring, while the other signal was located at the end of the other centriole (Fig. 1*B*, Top View, WT), indicating that SSNA1 localizes to the distal end of both mother and daughter centrioles. The side view further confirmed that SSNA1 localized to the distal end with CEP164 (Fig. 1*B*, Side View, WT), establishing SSNA1 as a distal centriolar protein.

### SSNA1 is present at the basal bodies of both primary and motile cilia

Since we detected SSNA1 at centrioles in dividing cells and at the basal bodies of primary cilia in serum-starved cells, we sought to determine whether SSNA1 is also present at the basal bodies of motile cilia (Fig. 2 *A* and *B*). To address this question, we used human nasal epithelial cells (HNECs). Stem cells were extracted from the nasal brushing of a healthy donor (HNEC0268) and then differentiated using air-liquid interface cultures to generate multicilia (57). Immunostaining with antibodies against SSNA1 and the basal body/SDA marker ODF2 revealed that SSNA1 was present and colocalized with ODF2 at the basal bodies of multiciliated cells (Fig. 2*A*).

**Fig. 2:**
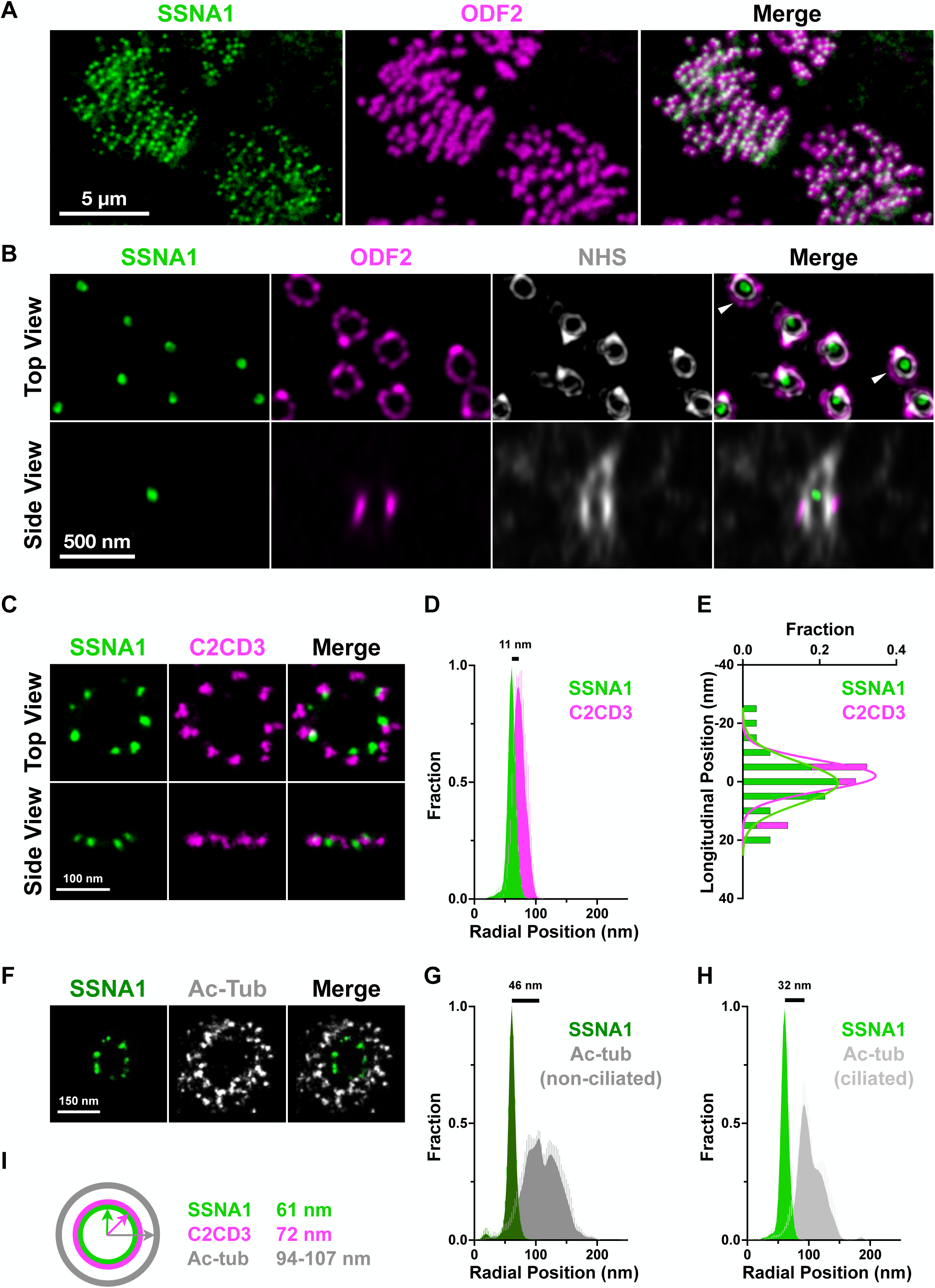
SSNA1 is present in primary and motile cilia and is arranged in a 9-fold ring-like pattern in the distal centriolar lumen. **A,** SSNA1 is present at the basal bodies of motile cilia. Stem cells were extracted from the nasal brushes of the HNEC0268 donor and then differentiated using air-liquid interface cultures to generate multiciliated cells. Immunostaining with antibodies against SSNA1 and the basal body/SDA maker ODF2 revealed that SSNA1 was present at the basal bodies of motile cilia. Bar, 5 µm. **B,** SSNA1 is localized to the distal lumen of the basal bodies. Representative TREx-confocal images of motile cilia in HNEC0268 cells. Cells were stained for SSNA1 and ODF2, along with a fluorophore-conjugated NHS ester to provide an ultrastructural context of the basal bodies. Top View: SSNA1 specifically localized inside the distal lumen of basal bodies in motile cilia. Arrowheads: Luminal SSNA1 does not directly contact the centriolar wall. Side View: SSNA1 localized more distally than the SDA protein ODF2. Expansion factor: 7.5-8x. Bar, 500 nm. **C,** Representative ExM-dSTROM images of centrioles in RPE-1 cells. Cells were stained with C2CD3 and SSNA1. SSNA1 is arranged in a ring-like pattern with near 9-fold symmetry in the distal lumen, and positioned adjacent to but more interior than C2CD3. Longitudinally, SSNA1 and C2CD3 had similar vertical positioning with minimal offset. Bar, 100 nm. **D,** Radial distribution analysis of SSNA1 and C2CD3. SSNA1 is more internally localized than C2CD3, with average radii measured at 60.9 nm for SSNA1 and 71.8 nm for C2CD3. n = 5 centrioles. **E,** Longitudinal distribution analysis of SSNA1 and C2CD3. Longitudinally, SSNA1 and C2CD3 had similar vertical positioning. **F,** Representative ExM-dSTROM images of centrioles in RPE-1 cells. Cells were stained for acetylated tubulin and SSNA1. SSNA1 is localized in the distal lumen distant from the centriolar microtubules. Bar, 150 nm. **G,** Radial distribution analysis of SSNA1 and acetylated tubulin in non-ciliated RPE-1 cells. The innermost peak radius of centriolar microtubules is 106.7 nm, approximately 46 nm away from SSNA1. n = 5 centrioles. **H,** Radial distribution analysis of SSNA1 and acetylated tubulin in ciliated RPE-1 cells. The innermost peak radius of centriolar microtubules is 93.2 nm, approximately 32 nm away from SSNA1. n = 5 centrioles. **I,** Schematic diagram of SSNA1 (60.9 nm) and C2CD3 (71.8 nm) radial positions within the distal lumen away from centriolar microtubules (93.2-106.7 nm). SSNA1 maintained a fairly consistent radial distribution in both ciliated and non-ciliated cells, regardless of ciliation-induced changes in microtubule arrangement.

### SSNA1 localizes to the distal lumen of centrioles

To define the precise localization of SSNA1 within the basal bodies of motile cilia, we performed Ten-fold Robust Expansion (TREx) (58) microscopy (Fig. 2*B*). Cells were stained for SSNA1 and ODF2, along with a fluorophore-conjugated N-hydroxysuccinimide (NHS) ester to provide an ultrastructural context of the basal bodies (59, 60). TREx-confocal imaging revealed that SSNA1 is specifically localized to the distal lumen of basal bodies in motile cilia (Fig. 2*B*, Top View) and longitudinally more distal than the SDA protein ODF2 (Fig. 2*B*, Side View). Notably, luminal SSNA1 does not directly contact the centriolar wall of the basal bodies. This is also clearly visible in some cases in the top view where the cilia are oriented at right angles (Fig. 2*B*, arrowheads), indicating that SSNA1 is located in the very center of the distal lumen. Despite the precise localization of SSNA1 to the distal lumen, the signal remained as a single isolated puncta at this resolution.

To further resolve the molecular arrangement of SSNA1 within the distal lumen, we applied ExM combined with direct stochastic optical reconstruction microscopy (dSTORM) to the expanded centrioles of proliferating and ciliated RPE-1 cells (Fig. 2 *C*-*I*). This advanced two-color single-molecule localization microscopy has allowed us to resolve the complex architecture of centriolar DA proteins at near molecular resolution (34, 36). Our recent work has shown that C2CD3, although located on the luminal side of the distal centriole, serves as an essential molecular hub organizing DA proteins outside the centriolar wall (36). Given the presence of SSNA1 in the distal lumen, we sought to define its precise spatial relationship with C2CD3 (Fig. 2 *C*-*E*). Using ExM-dSTORM, we observed SSNA1 as distinct puncta arranged in a near-9-fold symmetrical ring configuration and positioned adjacent to C2CD3 but slightly closer to the lumen center (Fig. 2*C*). Radial distribution analysis revealed that SSNA1 is more internally localized than C2CD3, with average radii of 71.8 nm for C2CD3 and 60.9 nm for SSNA1 (Fig. 2 *D* and *I*). Longitudinally, C2CD3 and SSNA1 show similar vertical positioning with minimal offset (Fig. 2*E*). The striking ring-like pattern and 9-fold symmetry of C2CD3 and SSNA1 is highly reminiscent of the distal ring density recently reported using cryo-electron tomography (cryo-ET) (61), suggesting that C2CD3 and SSNA1 may be part of the ordered assembly in the distal lumen of centrioles.

Furthermore, co-staining with acetylated tubulin revealed that SSNA1 resides relatively distant from centriolar microtubules (33-46 nm apart) (Fig. 2 *F*-*I*). In non-ciliated cells, centriolar microtubules present a broader radial distribution, with an innermost peak radius of 106.7 nm (Fig. 2 *G* and *I*). In ciliated cells, due to axoneme formation, the distal lumen narrows to a radius of 93.2 nm (Fig. 2 *H* and *I*). Importantly, SSNA1 maintains a fairly consistent radial distribution in both ciliated and non-ciliated cells, regardless of ciliation-induced changes in microtubule arrangement.

### SSNA1 does not associate with microtubules

SSNA1 has been reported as a microtubule remodeling factor capable of branching microtubules *in vitro* (53, 54). In contrast to another single-molecule TIRF study in which no branching was observed, purified His-SSNA1 was shown to stabilize microtubules and accumulate at the ends of growing microtubules and at sites of lattice damage (55). Despite these reported associations with microtubules *in vitro*, our investigation of its cellular localization indicates that SSNA1 is not present at the midbody or ciliary axoneme and has no direct contact with centriolar microtubules. The lack of microtubule-associated patterns in cells, together with the reported mismatch in axial periodicity between tubulin dimers (8 nm) (62, 63) and SSNA1 fibrils (11 nm) (53) *in vitro*, raised the question whether SSNA1 truly binds microtubules.

To determine whether SSNA1 directly binds microtubules *in vitro*, we purified full-length human SSNA1 and performed a microtubule co-pelleting assay using GST as a negative control and GM130 N74-GST, previously shown to bind microtubules *in vitro* (64), as a positive control. We bacterially expressed and affinity purified four recombinant SSNA1 fusion proteins with either an N- or C-terminal GST or His tag (GST-SSNA1, SSNA1-GST, His-SSNA1, and SSNA1-His) (all His tags used in this study are His6) (Fig. 3*A*). To allow tag removal, a protease cleavage site (3C for GST and TEV for His) was inserted between the tag and SSNA1. Tag-free SSNA1 was obtained by cleaving GST-SSNA1 with 3C protease followed by size exclusion chromatography (SEC) purification (Fig. S4 *A* and *B*, 1-119/FL and WT). The resulting tag-cleaved SSNA1 retained only two additional amino acids (Gly-Pro) at the N-terminus, closely resembling native SSNA1 *in vivo*.

**Fig. 3:**
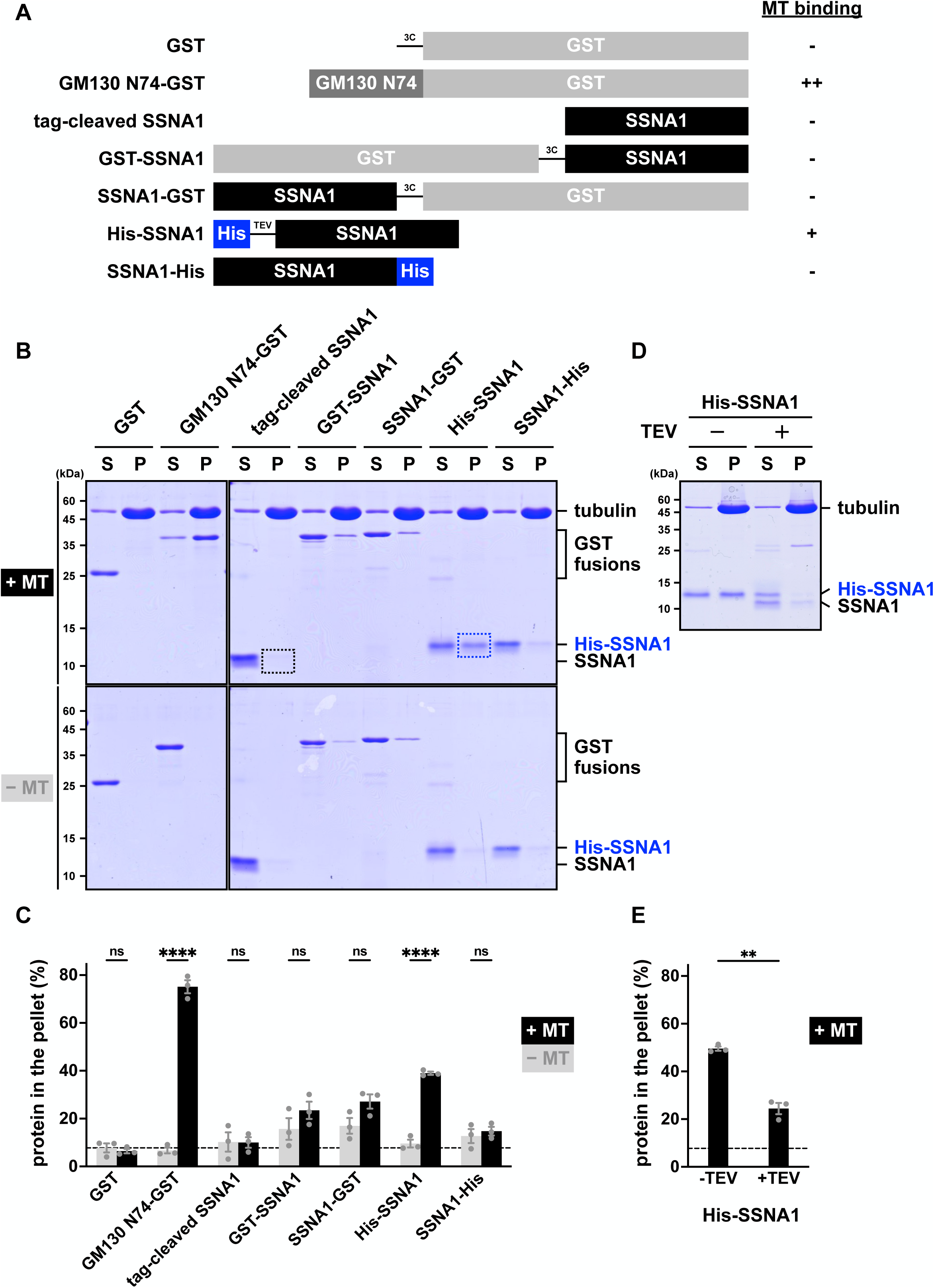
SSNA1 is not a microtubule-associated protein. **A,** Left: schematic diagram of SSNA1 constructs used to assess microtubule binding *in vitro*. GST and GM130 N74-GST served as negative and positive controls, respectively. Recombinant SSNA1 with N- or C-terminal GST or His tags was bacterially expressed and affinity purified. Protease cleavage sites (3C or TEV) were incorporated to facilitate tag removal, as illustrated. Tag-cleaved SSNA1 was derived from GST-SSNA1. Right: summary of microtubule binding in **B**. **B,** Microtubule co-pelleting assay. Proteins incubated with (+MT) or without (-MT) microtubules were subjected to ultracentrifugation to separate supernatant (S) from pellet (P) fractions. None of the recombinant SSNA1 proteins bound microtubules and they remained in the supernatant, except for His-SSNA1, which exhibited a partial, non-specific microtubule association. **C,** Quantification of band intensities from Coomassie blue-stained gels in **B**. Protein in the pellet (%) was calculated as P/(S+P). Error bars represent the mean ± SEM. N=3. ****p < 0.0001; ns: not significant; two-way ANOVA. **D,** His-SSNA1 was treated with (+) or without (-) TEV protease prior to ultracentrifugation. His-SSNA1 dissociated from microtubules and shifted to the supernatant fraction upon His tag removal. **E,** Quantification of band intensities from Coomassie blue-stained gels in **D**. Protein in the pellet (%) was calculated as P/(S+P). Error bars represent the mean ± SEM. N=3. **p < 0.01; paired t test.

Next, we performed the microtubule co-pelleting assay with seven purified proteins: GST, GM130 N74-GST, tag-cleaved SSNA1, GST-SSNA1, SSNA1-GST, His-SSNA1, and SSNA1-His (Fig. 3*B*). As controls, all proteins were subjected to ultracentrifugation in the absence of preformed microtubules (Fig. 3*B*, -MT). Notably, trace amounts of GST-SSNA1 and SSNA1-GST appeared in the pellet, likely due to protein aggregation induced by GST dimerization and the inherent tendency of SSNA1 to oligomerize (Figs. 4 and S4). Nevertheless, despite their strong propensity to self-assemble into fibrils (Fig. 4*C*), all SSNA1 proteins remained predominantly soluble and were detected in the supernatant.

**Fig. 4:**
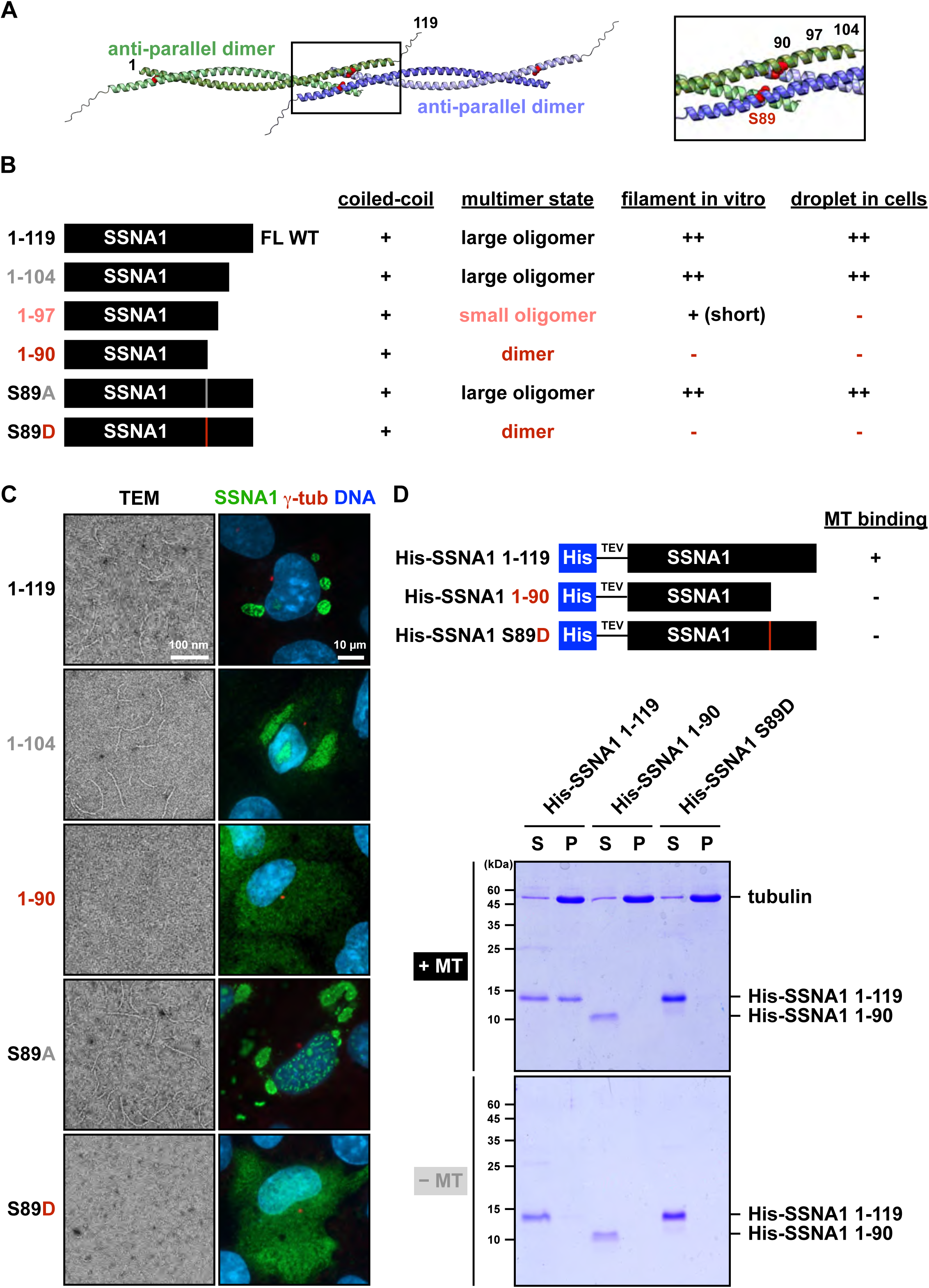
SSNA1 oligomerization induces nonspecific binding of His-SSNA1 to microtubules. **A,** Left: the predicted structure of the SSNA1 tetramer modeled by ColabFold (65). Two SSNA1 monomers form an antiparallel dimer through coiled-coil interactions, and two dimers further interact through C-terminal α-helices to form a tetramer, likely underlying the ability of SSNA1 to self-oligomerize. The C-terminal α-helices and residue S89 play essential roles in mediating interdimer association and oligomerization. Right: magnification of the inset on the left, showing relative positions of the key residues used to make oligomerization-deficient mutants, including residues 104, 97, 90 for truncation and S89 in red. **B,** Left: schematic diagram of SSNA1 constructs. Right: summary of their properties, including the ability to form coiled-coils (Fig. S5), the multimeric state *in vitro* (Fig. S4), the ability to form filaments *in vitro* (Fig. S6), and the ability to form droplets in cells (Fig. S8). For *in vitro* assays, all constructs were bacterially expressed and affinity purified as GST-SSNA1, followed by 3C protease cleavage and SEC purification. For the *in vivo* droplet formation assay, all constructs were expressed without any tags in SSNA1 KO RPE-1 cells. SSNA1 1-119 (full-length WT), C-terminal truncated mutants (1–104, 1–97, 1–90) and S89 point mutants (S89A, S89D) are presented. Two oligomerization-deficient mutants, SSNA1 1-90 and SSNA1 S89D, were identified. **C,** Left: representative negative-staining TEM images of recombinant SSNA1. SSNA1 1-90 and S89D failed to form filaments *in vitro*. Bar, 100 nm. Right: representative immunofluorescence images of overexpressed SSNA1 in SSNA1 KO RPE-1 cells. Cells were stained for SSNA1, γ-tubulin and DNA. SSNA1 1-90 and S89D did not form droplets *in vivo*. Bar, 10 µm. **D,** Microtubule co-pelleting assay. SSNA1 1-119 (full-length WT) and two oligomerization-deficient mutants 1-90 and S89D were purified as His-SSNA1, incubated with (+MT) or without (-MT) microtubules, followed by ultracentrifugation to separate supernatant (S) from pellet (P) fractions. While His-SSNA1 1-119 exhibited partial microtubule binding, the two oligomerization-deficient mutants His-SSNA1 1-90 and His-SSNA1 S89D lost the association with microtubules, indicating that SSNA1 oligomerization induces non-specific binding of SSNA1 fibrils to microtubules.

To assess direct microtubule association, we added preformed taxol-stabilized microtubules to the purified proteins and repeated the ultracentrifugation (Fig. 3*B*, +MT). As expected, GST remained in the supernatant, whereas GM130 N74-GST was found in the pellet, validating the efficacy of the assay. In addition, both GST-SSNA1 and SSNA1-GST remained in the supernatant, with trace amounts detected in the pellet comparable to controls without microtubules. Similarly, SSNA1-His remained in the supernatant, confirming a lack of microtubule binding. Most importantly, tag-cleaved SSNA1 (Fig. 3*B*, black box) exhibited the same behavior as SSNA1-His, reinforcing that SSNA1 does not bind microtubules *in vitro*. In contrast, His-SSNA1 exhibited partial microtubule binding (Fig. 3*B*, blue box, and 3*D*, -TEV), with approximately 40-50% of the protein bound to microtubules and shifted to the pellet (Fig. 3 *C* and *E*). These data indicate that previous studies (53, 55) using His-SSNA1 may have observed non-specific binding. To test this possibility, we treated His-SSNA1 with TEV protease for 10 minutes before performing the microtubule co-pelleting assay (Fig. 3 *D* and *E*). After treatment, both cleaved and uncleaved SSNA1 proteins were found in the supernatant, indicating that removal of the His tag from SSNA1 was sufficient to eliminate non-specific binding to microtubules.

### Identification and characterization of SSNA1 oligomerization-deficient mutants

Since tag-cleaved SSNA1 does not bind microtubules *in vitro*, we postulated that the reported longitudinal association of His-SSNA1 fibrils along microtubules (53) might result from weak binding of the charged His tag, which is further enhanced by SSNA1 oligomerization. To investigate this possibility, we designed SSNA1 mutants deficient in self-oligomerization, guided by the predicted structure of the SSNA1 tetramer modeled by ColabFold (65) (Fig. 4*A*). Structurally, SSNA1 is composed of a predominantly α-helical domain with multiple heptad repeats that drive coiled-coil formation (45, 48, 52–54), followed by an intrinsic disordered region (IDR) at the C-terminus. Structural predictions indicated that two SSNA1 monomers form an antiparallel dimer through coiled-coil interactions, with two dimers further interacting via their C-terminal α-helical regions to form a tetramer, likely underlying its ability to self-oligomerize.

We hypothesized that the C-terminal α-helices might be essential for interdimer associations and thus oligomerization. To test this possibility, we sequentially truncated the C-terminus, reducing the protein length from residues 1-119 (full-length, FL) to 1-104 (removing the IDR, ΔIDR), 1-97 (removing the IDR plus 7 additional amino acids), and 1-90 (removing the IDR plus 14 additional amino acids) (Fig. 4*B*). Each truncated protein was affinity purified as GST-SSNA1, with the GST tag subsequently cleaved and further purified by SEC (Fig. S4*A*). To ensure that the truncations did not alter the secondary structure or coiled-coil dimerization of SSNA1, all purified tag-cleaved proteins were subjected to circular dichroism (CD) analysis (Fig. S5). CD spectra for the full-length and truncated proteins displayed characteristic α-helical profiles, with a positive band ∼190 nm and two negative minima at 208 nm and 222 nm (Fig. S5*A*). The molar ellipticity ratio at 222 nm to 208 nm (222/208) was greater than one for all proteins (Fig. S5*C*), indicating that the coiled-coil structure remained intact for all truncated variants (Fig. 4*B*).

Further SEC analysis revealed that the full-length protein (residues 1-119) eluted from the void volume and was widely distributed across almost all fractions (Fig. S4*A*), indicating that SSNA1 self-assembles into heterogeneous higher-order oligomers. Consistent with previous reports (48, 49, 53), negative staining transmission electron microscopy (TEM) (Figs. 4*C* and S6*A*) showed these oligomers to be long filaments averaging 55 nm in length and 4.3 nm in diameter (Fig. S6*C*). The 1-104 (ΔIDR) truncation variant also eluted from the void volume, but it exhibited a more defined size, indicating that removal of the IDR may promote more uniform oligomerization. Our TEM analysis also revealed that the 1-104 variant generated long fibrils comparable to those formed by full-length SSNA1. In contrast, the 1-97 truncation variant eluted in later fractions and appeared as very short filaments in TEM images. The 1-90 truncation variant eluted in much later fractions and fibrils were not detected by TEM. By multi-angle light scattering (MALS) analysis, the molecular weight of SSNA1 decreased from an estimated 1 MDa for the full-length oligomers to 18.6 kDa for the 1-90 truncation variant, consistent with a dimer (Figs. 4*B* and S4 *C* and *D*).

Next, we examined the effects of SSNA1 overexpression in cells. To this end, we transfected a construct expressing untagged full-length human SSNA1 into four cell lines, including HEK, HeLa, as well as WT and SSNA1 KO RPE-1 cells. Overexpressed SSNA1 self-associated and formed droplet-like structures in all four cell lines (Fig. S7*A*). To rule out the possibility that SSNA1 self-association was initiated by or dependent on endogenous SSNA1, we performed all further overexpression experiments in the SSNA1 KO RPE-1 cells. We observed a wide range of droplet sizes depending on SSNA1 expression levels (Fig. S7*B*). Although SSNA1 at low expression levels occasionally appeared as small dots near the centrosomal marker γ-tubulin, the majority of overexpressed SSNA1 did not localize to centrosomes. Instead, the self-assembled droplets formed in both the nucleus and cytoplasm (Fig. S7*C*), evidencing a strong propensity for overexpressed SSNA1 to self-associate rather than target to centrosomes. Quantification of droplet localization revealed that 8.0% of the SSNA1-overexpressing cells solely hosted droplets in the nucleus, 16.6% only in the cytoplasm, and 75.4% had droplets in both compartments. In contrast to the numerous small puncta observed throughout the nucleus reported in the first study that identified SSNA1 as a nuclear antigen (45), we observed fewer but larger droplets. Confocal imaging confirmed that, when present in the nucleus, these larger droplets were often Hoechst negative and occupied a DNA-free space inside the nucleus. The strong tendency of SSNA1 to self-associate seems to push DNA away and exclude it from co-assembly (*e.g*., Figs. 5*A*, S8 and S9). Although endogenous SSNA1 is not a nuclear resident protein under physiological conditions, the presence of nuclear droplets suggests that highly expressed SSNA1 can enter the nucleus due to its small size (a dimer is 28 kDa, below the free diffusion threshold of the nuclear pore complex).

**Fig. 5:**
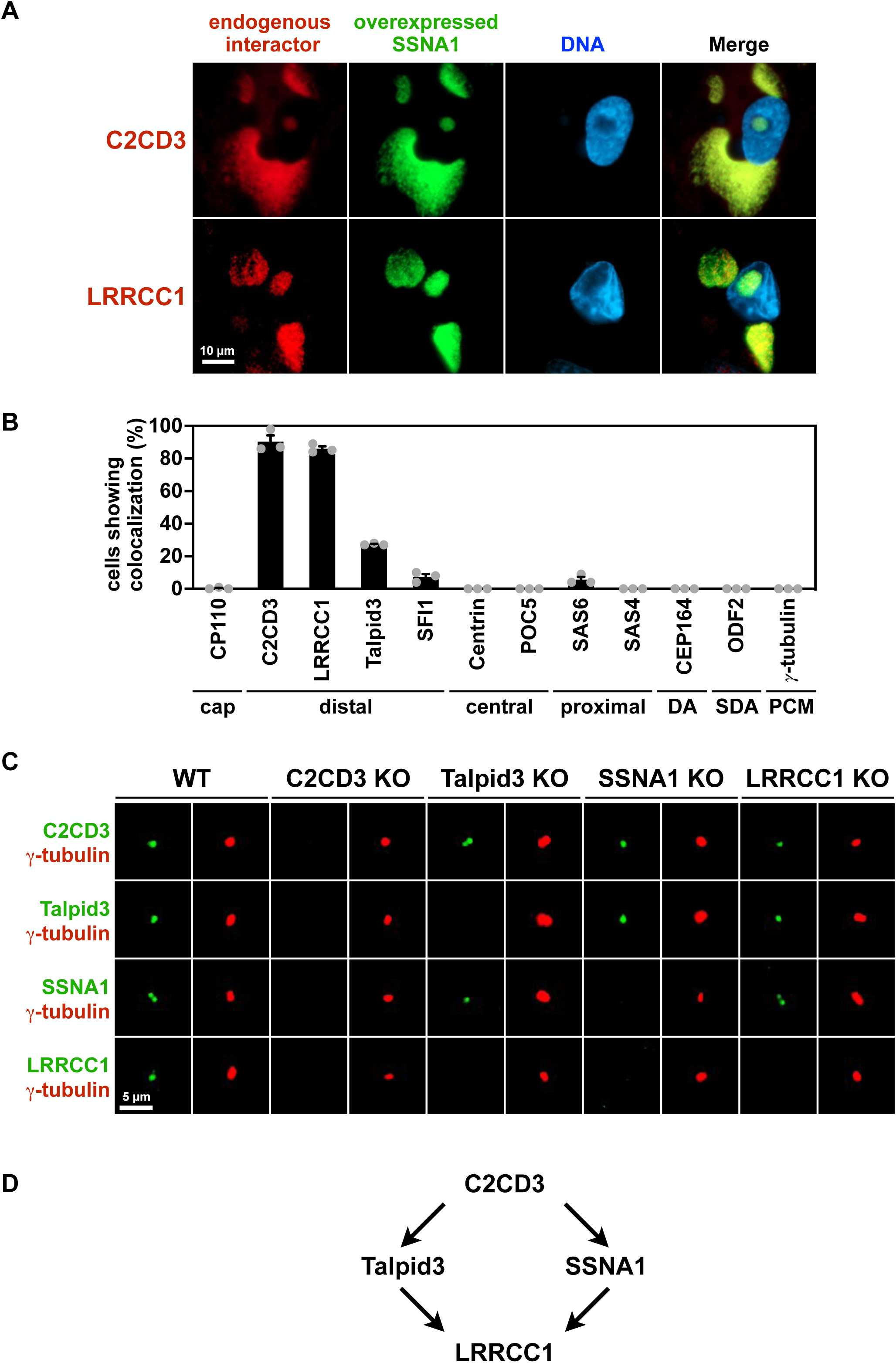
SSNA1 organizes a hierarchical assembly pathway comprising the C2CD3-SSNA1-LRRCC1 axis at the distal lumen. **A,** SSNA1 interacts with C2CD3 and LRRCC1. SSNA1 KO RPE-1 cells expressing untagged full-length SSNA1 were stained with antibodies against SSNA1 and candidate interacting proteins. Overexpressed SSNA1 dissociated its endogenous binding partners C2CD3 and LRRCC1 from the centrosomes and redirected them into its self-assembled droplets. Bar, 10 µm. **B,** Quantification of colocalization in **A**. The percentage of SSNA1-expressing cells showing colocalization of candidate proteins with SSNA1-positive droplets was scored. Representative markers of different centrosomal regions, including the axoneme cap (CP110), distal lumen (C2CD3, LRRCC1, SFI1), central lumen (Centrin, POC5), proximal end/procentriole (SAS6, SAS4), DA (CEP164), SDA (ODF2), and PCM (γ-tubulin), were stained and quantified. Only two distal luminal proteins, C2CD3 and LRRCC1, colocalized with SSNA1. Error bars represent the mean ± SEM. N=3, n > 100 cells per experiment. **C,** Centriolar targeting of C2CD3, Talpid3, SSNA1, and LRRCC1 in RPE-1 WT and individual KO cells. Cells were stained with antibodies against C2CD3, Talpid3, SSNA1, and LRRCC1, along with γ-tubulin to label centrosomes. In WT cells, all proteins were detected at centrosomes. In C2CD3 KO cells, signals for Talpid3, SSNA1, and LRRCC1 were completely absent, indicating that C2CD3 functions as the primary recruiter. Taplid3 KO cells retained the centrosomal localization of C2CD3 and SSNA1, but the LRRCC1 signal was lost, suggesting that Talpid3 regulates LRRCC1 recruitment. Similarly, in SSNA1 KO cells, the centrosomal localization of C2CD3 and Talpid3 persisted while LRRCC1 targeting was disrupted, indicating that SSNA1 is also required for LRRCC1 recruitment, but through a Talpid3-independent pathway. The centrosomal localization of C2CD3, Talpid3 and SSNA1 remained intact in LRRCC1 KO cells, placing LRRCC1 as the most downstream component in this recruitment hierarchy. Bar, 5 µm. **D,** Schematic diagram of the hierarchical recruitment of C2CD3, Talpid3, SSNA1, and LRRCC1. C2CD3 serves as the most upstream recruiter, whereas Talpid3 and SSNA1 act as intermediate regulators to facilitate LRRCC1 localization.

Next, we expressed untagged SSNA1, including the full-length (1–119) and truncated variants (1-104, 1-97, and 1-90) in SSNA1 KO cells (Figs. 4*C* and S8). Overexpressed untagged SSNA1 1-119 and 1-104 formed droplets, whereas 1-97 and 1-90 variants were dispersed throughout the cells. Together with our TEM analysis, these data indicate that the ability of SSNA1 to form filaments *in vitro* is correlated with its capacity to form droplets in cells, with both serving as proxies for self-oligomerization. Thus, the SSNA1 1-90 variant represents an oligomerization-deficient mutant.

To rule out the possibility that truncation removed key motifs necessary not only for oligomerization but also for microtubule binding, we sought to generate a second oligomerization-deficient mutant harboring a single point mutation yet retaining the full protein length. Based on the ColabFold-predicted structure of the SSNA1 tetramer, we identified serine (S) residue 89 as a critical residue for oligomerization because it faces another S89 residue from the other dimer at the interdimer interface (Fig. 4*A*). We reasoned that substitution of S89 with negatively charged aspartic acid (D) would introduce electrostatic repulsion between the two mutated residues, disrupt the dimer interface, and prevent oligomerization. As a control, we also substituted S89 with the non-polar residue alanine (A). We purified the S89A and S89D point mutants and confirmed their coiled-coil configuration by CD analysis (Fig. S5 *B* and *D*). Whereas S89A eluted similarly to the WT protein in SEC, S89D eluted in much later fractions and slightly earlier than SSNA1 1-90 (Fig. S4*B*). SEC-MALS analysis revealed that S89D had a molecular weight of 31.1 kDa, consistent with a dimer (Fig. S4 *C* and *D*). TEM analysis showed that S89A formed long filaments similar to WT, whereas S89D did not assemble into fibrils *in vitro* (Figs. 4*C* and S6*B*). Consistently, S89A formed droplets, whereas S89D was dispersed throughout the cells (Figs. 4*C* and S8). Thus, SSNA1 S89D represents a second oligomerization-deficient mutant.

### SSNA1 oligomerization contributes to non-specific binding of His-SSNA1 to microtubules

To test our hypothesis that SSNA1 oligomerization promotes non-specific binding of His-SSNA1 to microtubules, we affinity purified the two newly identified oligomerization-deficient mutants, *i.e.*, 1-90 and S89D, in the form of His-SSNA1, and repeated the microtubule co-pelleting assay (Fig. 4*D*). As protein-only controls, the three His-SSNA1 proteins (His-SSNA1 1-119/WT, His-SSNA1 1-90 and His-SSNA1 S89D) remained soluble. Upon addition of preformed microtubules, His-SSNA1 1-119/WT was partially shifted into the pellet, as described above. In contrast, the two oligomerization-deficient mutants, His-SSNA1 1-90 and His-SSNA1 S89D, remained exclusively in the supernatant (Fig. 4*D*). This result indicates that disruption of SSNA1 oligomerization and filament formation effectively prevents non-specific binding to microtubules. Taken together, our results demonstrate that the previously observed longitudinal association of His-SSNA1 filaments along microtubules is caused by non-specific binding of the His tag exacerbated by SSNA1 oligomerization. These results, together with our *in vivo* findings that SSNA1 does not localize to the midbody, ciliary axoneme or centriolar microtubules, support a revision of the role of SSNA1 and indicates that it does not nucleate, stabilize or branch microtubules as previously proposed.

### SSNA1 interacts with C2CD3 and LLRCC1 at the distal lumen of centrioles

The lack of binding of SSNA1 to centriolar microtubules (Figs. 2 *B* and 3), together with its close proximity to C2CD3 in the distal lumen (Fig. 2 *C*-*E*), prompted us to investigate possible direct interactions between SSNA1 and C2CD3. We also examined the binding of SSNA1 to other proteins that either associate with C2CD3 (*e.g*., Talpid3 (66)) or localize to the distal lumen of centrioles (*e.g*., LRRCC1 (43) and SFI1 (44)).

To test for interactions, we took advantage of the cellular droplet formation assay described above (Figs. 5 *A* and *B* and S9). Overexpression of untagged full-length human SSNA1 in SSNA1 KO RPE-1 cells resulted in droplet self-assembly. The formation of SSNA1 droplets may incorporate its native binding partners and sequester them from the centrioles. Therefore, we stained the transfected cells with the SSNA1 A6 antibody to locate droplets together with antibodies against candidate proteins (C2CD3, LRRCC1, Talpid3, and SFI1). We then evaluated the percentage of SSNA1-expressing cells that showed colocalization of candidate proteins with SSNA1 droplets (Fig. 5*B*). As a control, we analyzed proteins representing different regions of centrosomes, including distal cap (CP110), central lumen (Centrin, POC5), proximal end/procentriole (SAS6, SAS4), DA (CEP164), SDA (ODF2), and PCM (γ-tubulin) (Figs. 5*B* and S9). All of these markers remain localized to centrosomes, indicating no interaction with SSNA1. In contrast, endogenous C2CD3 and LRRCC1 were dissociated from centrosomes and strongly colocalized with SSNA1 droplets (Fig. 5*A*), supporting that both C2CD3 and LRRCC1 interact with SSNA1 at the distal lumen of centrioles. Notably, SFI1 and Centrin, which form a complex in the distal lumen (44), did not overlap with SSNA1 droplets, suggesting that C2CD3/SSNA1/LRRCC1 and SFI1/Centrin may function independently in the distal lumen. Talpid3 appeared in SSNA1-positive droplets in a small percentage of cells, but with a very low degree of overlap (Fig. S9), which may be due to its binding to C2CD3.

### Hierarchical assembly of the C2CD3-SSNA1-LRRCC1 targeting axis

Given the strong relocation of endogenous C2CD3 and LRRCC1 from centrosomes to SSNA1 droplets, we determined whether these interactions occur under physiological conditions and whether their centriolar localizations are interdependent. To address these possibilities, we generated KO RPE-1 cells depleted of individual distal luminal proteins, including C2CD3, Talpid3 (which binds C2CD3, although likely not in the lumen (66)), and LRRCC1, but were unable to obtain KO clones for SFI1 (Fig. 5*C* and Supplementary Table 2). Western blotting analysis confirmed the absence of these proteins in total cell lysates of the respective KO cells (Fig. S10), and immunofluorescence further validated the loss of their centrosomal signals (Fig. 5*C*). These KO cell lines allow us to define the interdependencies and functional roles of distal luminal proteins in centriolar organization.

We then immunostained RPE-1 WT and individual KO cells with antibodies against C2CD3, Talpid3, SSNA1, and LRRCC1, as well as γ-tubulin to label centrosomes (Fig. 5*C*). All proteins were detected at centrosomes in WT cells. However, signals for Talpid3, SSNA1, and LRRCC1 were absent in C2CD3 KO cells, indicating that C2CD3 acts as an upstream recruiter of these proteins to distal centrioles. In contrast, C2CD3 and SSNA1, but not LRRCC1, localized to centrosomes in Taplid3 KO cells, suggesting that Talpid3 regulates the centriolar targeting of LRRCC1. Similarly, in SSNA1 KO cells, the centrosomal localization of C2CD3 and Talpid3 persisted, whereas LRRCC1 targeting was disrupted, implying that SSNA1 is also required for LRRCC1 recruitment, but through a Talpid3-independent pathway. Finally, the centrosomal localization of C2CD3, Talpid3 and SSNA1 remained intact in LRRCC1 KO cells, placing LRRCC1 as the most downstream component in this recruitment hierarchy (Fig. 5*D*). Notably, the absence of signals at centrosomes in KO cells was due to a failure of centriolar targeting rather than protein destabilization, as total protein levels were unchanged (Fig. S10). Taken together, our results delineate a hierarchical targeting axis for the recruitment of distal luminal proteins to centrioles. C2CD3 serves as an upstream recruiter, whereas Talpid3 and SSNA1 act as intermediate regulators to facilitate LRRCC1 localization (Fig. 5*D*). This network highlights the coordinated assembly of centriolar components essential for distal lumen organization.

### SSNA1 is not required for overall centriole architecture and appendage assembly

Depletion of C2CD3 has been shown to disrupt the recruitment of several DA proteins located outside the centriolar lumen (40), as well as the luminal proteins demonstrated in our study. Since SSNA1 functions downstream of C2CD3 in the distal targeting network, we investigated whether SSNA1 KO affects the organization of other centriolar regions. To evaluate this possibility, we examined representative markers for different regions of centrosomes as listed above. All of the proteins we tested remained localized to centrosomes in SSNA1 KO cells, indicating that SSNA1 is not required to maintain the overall architecture of centrioles or centrosomes (Fig. 6*A*).

**Fig. 6:**
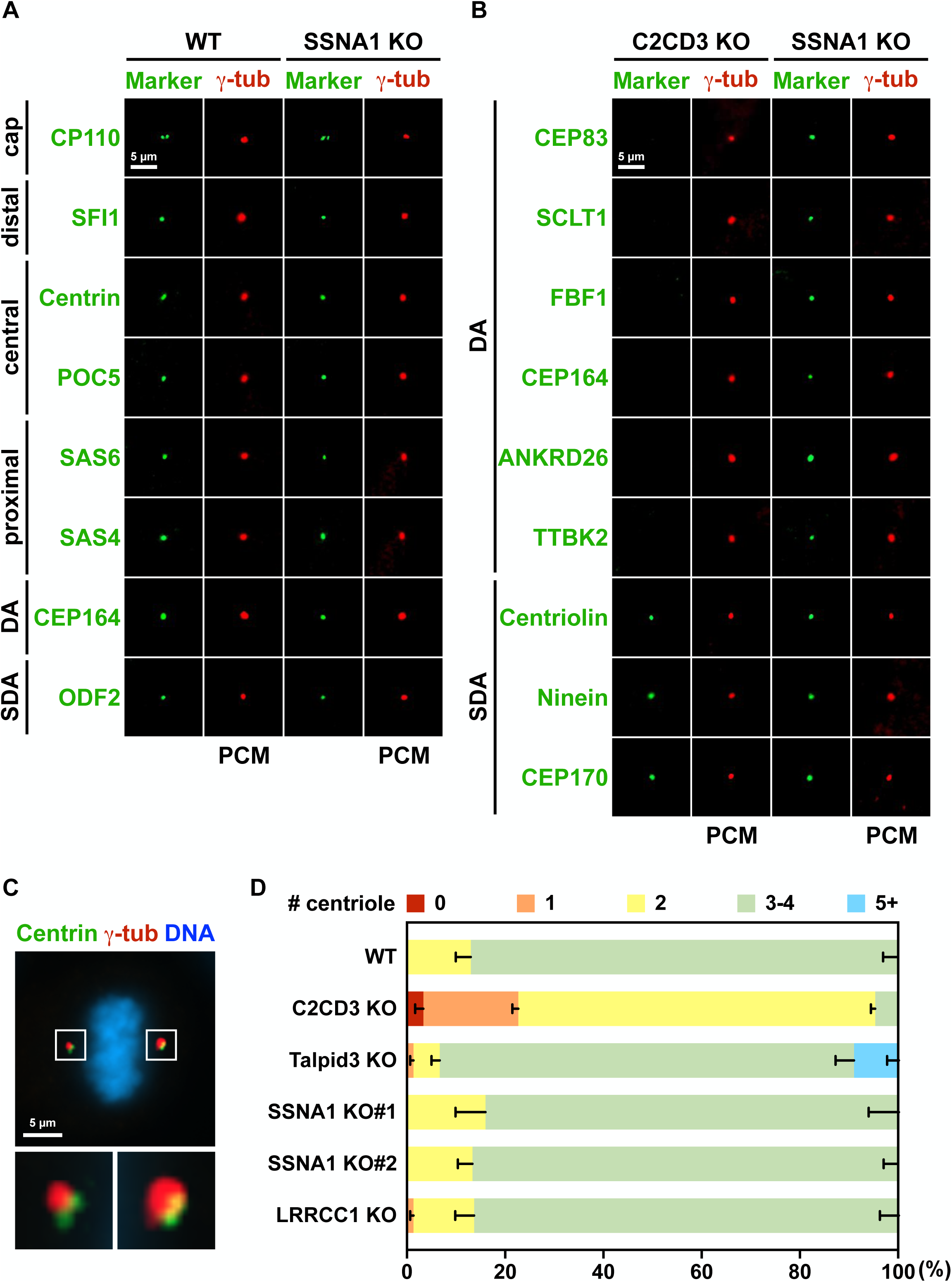
SSNA1 is not required for overall centriolar architecture or duplication. **A,** WT and SSNA1 KO RPE-1 cells were stained with antibodies against the representative markers of different centriolar regions, along with γ-tubulin to label centrosomes. All the markers tested, including CP110, SFI1, Centrin, POC5, SAS6, SAS4, CEP164, ODF2, and γ-tubulin, were correctly targeted in SSNA1 KO cells, indicating that SSNA1 is not required to maintain the overall architecture of centrioles or centrosomes. Bar, 5 µm. **B,** C2CD3 KO and SSNA1 KO RPE-1 cells were stained with antibodies against the representative markers of DA and SDA, along with γ-tubulin to label centrosomes. Despite the massive loss of DA targeting in C2CD3 KO cells, all the DA and SDA markers tested, including CEP83, SCLT1, FBF1, CEP164, ANKRD26, TTBK2, Centriolin, Ninein, CEP170, were correctly targeted in SSNA1 KO cells. Unlike C2CD3 KO, where DA assembly outside the lumen is severely disrupted, SSNA1 KO only affected distal luminal organization. Bar, 5 µm. **C,** RPE-1 cells were stained with antibodies against Centrin to label centrioles and γ-tubulin to label centrosomes. Only metaphase cells were used to score the centriole number per mitotic cell. A representative image of a metaphase cell showing two centrioles per mitotic centrosome at each pole, resulting in a total of 4 centrioles per mitotic cell. Bar, 5 µm. **D,** The number of centrioles in mitotic centrosomes of metaphase cells was assessed in WT and individual KO cells. In contrast to C2CD3 KO cells most of which contained only two centrioles per mitotic cell, SSNA1 KO and LRRCC1 KO cells had normal centriole numbers, indicating that SSNA1 is not required for centriole biogenesis. N=3, n = 40-50 metaphase cells per experiment.

Although located within the centriolar lumen, C2CD3 mediates the assembly of DA and possibly SDA outside the distal lumen (40, 66). Therefore, we tested whether the DA proteins that depend on C2CD3 for targeting, including CEP83, SCLT1, FBF1, CEP164, ANKRD26, and TTBK2, and the SDA proteins, including Centriolin, Ninein, and CEP170, are recruited to the centrioles in SSNA1 KO cells (Fig. 6*B*). Consistent with previous reports (40, 66), all of the DA proteins we tested lost their centriolar targeting in C2CD3 KO cells, but remained localized to centrosomes in SSNA1 KO cells. In contrast, the centrosomal localization of SDA proteins was unaffected in both C2CD3 KO and SSNA1 KO cells. These results indicate that C2CD3 is required for DA but not for SDA assembly and that, unlike C2CD3 KO cells in which DA assembly outside the lumen is severely disrupted, SSNA1 only affects organization within the distal lumen.

### SSNA1 is not required for centriole duplication or cell division

To investigate whether SSNA1 is involved in centriole duplication, we immunostained cells for Centrin to label centrioles and γ-tubulin to label centrosomes. We then counted the number of centrioles at metaphase (Fig. 6*C*) (67) in WT and KO cells (Fig. 6*D*). As expected, the majority of WT cells contained two centrioles per centrosome at each pole, resulting in a total of four centrioles per mitotic cell. However, the majority of C2CD3 KO cells contained only one Centrin-positive centriole at each pole, resulting in a total of two Centrin-positive centrioles per mitotic cell, consistent with the established role of C2CD3 in centriole elongation (68). In contrast, ∼10% of Talpid3 KO cells had more than five centrioles per mitotic cell, indicating a mild defect caused by either centriole overamplification or cytokinesis failure. Nevertheless, SSNA1 KO and LRRCC1 KO cells had a normal number of centrioles similar to WT, supporting the notion that distal luminal proteins are not involved in centriole biogenesis.

Previous studies have suggested that SSNA1 localizes to centrosomes and midbodies to regulate mitosis and cytokinesis (46, 51). However, as we demonstrated here, the reported midbody localization was due to antibody cross-reactivity (Fig. S3*A*). We also successfully knocked out SSNA1 in several cell lines, including human RPE-1 and HeLa cells (not shown), as well as mouse 3T3 and mIMCD3 cells (see below for KO generation). Their viability and proliferation indicate that SSNA1 is not essential for cell division in mammalian cells, similar to SSNA1 KO in protozoa (48, 49). Furthermore, we found that the doubling time of two SSNA1 KO clones was comparable to WT, suggesting that SSNA1 does not play a significant role in regulating cell cycle progression or cell division. Instead, its primary function may involve pathways unrelated to cell proliferation, such as signaling, differentiation, or specialized cellular processes.

### SSNA1 regulates primary ciliogenesis of both intracellular and extracellular pathways

In this context, C2CD3 and LRRCC1, the upstream and downstream interactors of SSNA1, have been shown to regulate ciliogenesis (40, 42, 43). C2CD3 functions in vertebrate development and its mutations lead to a variety of ciliopathies, including orofaciodigital syndrome, polydactyly, and skeletal dysplasia (68–70). As shown here, SSNA1 is present in the basal bodies of both primary and motile cilia (Figs. 1 and 2). Therefore, we investigated whether depletion of SSNA1 or other distal proteins, including C2CD3, Talpid3, and LRRCC1, affects primary ciliogenesis in RPE-1 cells. We induced primary cilia in WT and KO cells by serum starvation and stained for Arl13b to label cilia and for the SDA marker Ninein to label the mother centriole (Fig. 7*A*), as DA proteins were no longer recruited in C2CD3 KO cells. The percentage of ciliated cells was then analyzed (Fig. 7*B*). In WT cells, ∼70% of the cells formed cilia 24 hours after serum starvation. In contrast, both SSNA1 KO cell lines showed a significant decrease in the proportion of ciliated cells to 22% for KO#1 and 20% for KO#2 (Fig. 7*B*), supporting that SSNA1 is required for efficient primary ciliogenesis in RPE-1 cells. As expected, C2CD3 KO and Talpid3 KO cells with severely disrupted DA assembled essentially no cilia, whereas LRRCC1 KO cells also showed a significant albeit slightly moderate reduction in cilia ratio, comparable to that induced by LRRCC1 knockdown (43). Interestingly, the lower cilia ratio observed in SSNA1 KO than in LRRCC1 KO further suggests that the more severe ciliogenesis defects in SSNA1 KO cells are not entirely caused by LRRCC1 loss and that other factors downstream of SSNA1 must contribute. Thus, a complete targeting and functional network at the distal lumen awaits further elucidation.

**Fig. 7:**
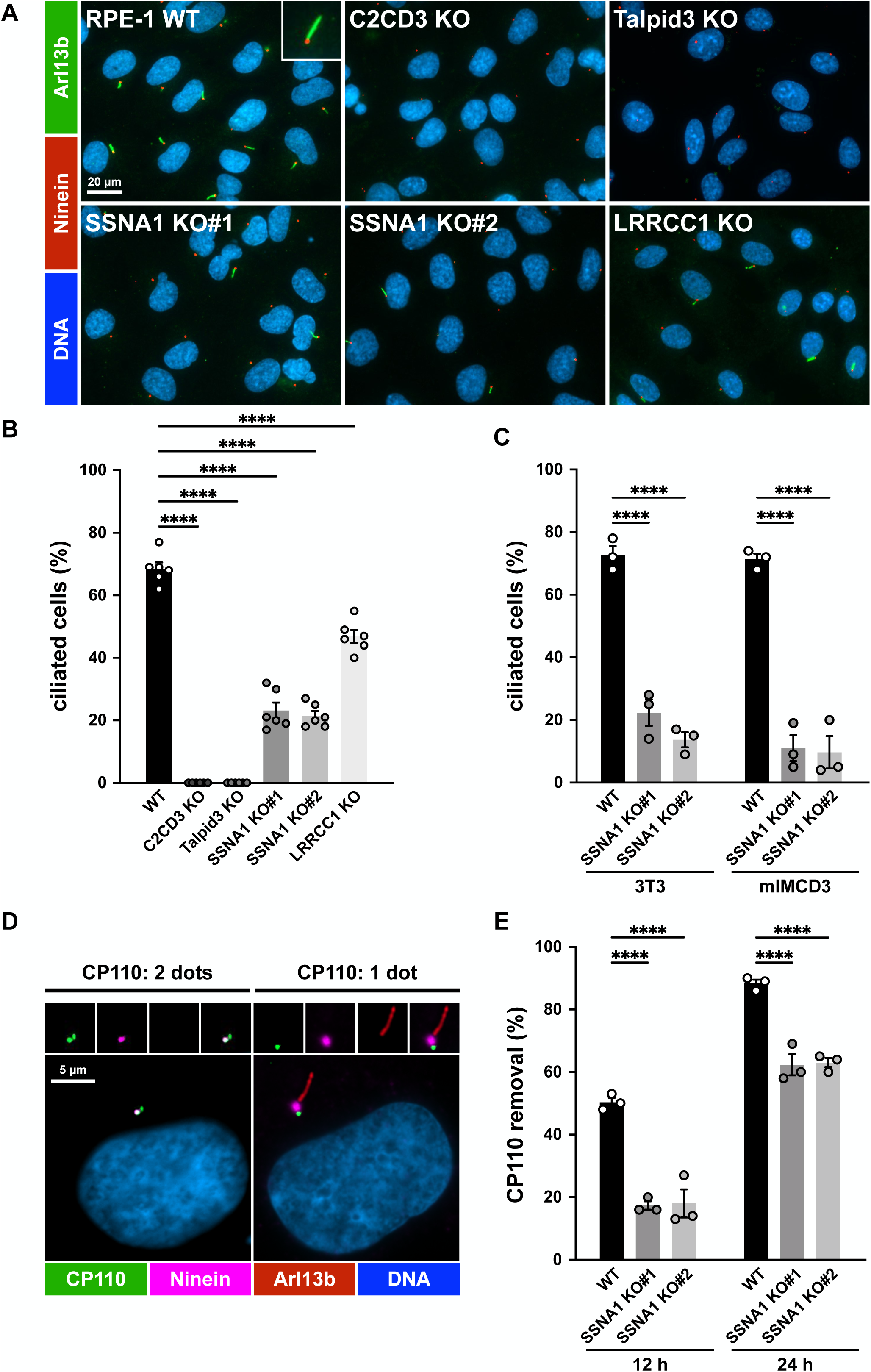
SSNA1 promotes efficient ciliogenesis of both intracellular and extracellular pathways by facilitating CP110 removal. **A,** Representative images of RPE-1 WT and individual KO cells after 24-hour serum starvation. Cells were stained for the ciliary membrane marker Arl13b (green), the SDA marker Ninein (red), and DNA (blue). Bar, 20 µm. A typical cilium is shown in the inset. **B,** To assess cilia assembly, RPE-1 WT and individual KO cells were induced to form primary cilia by serum starvation for 24 hours. The percentage of ciliated cells was quantified. SSNA1 KO resulted in significantly decreased ratios of ciliated cells. Error bars represent the mean ± SEM. N=6. ****p < 0.0001; one-way ANOVA. **C,** SSNA1 in 3T3 cells (that use the intracellular pathway) and in mIMCD3 cells (that use the extracellular pathway) were knocked out. WT and SSNA1 KO cells were induced to form primary cilia by serum starvation for 24 hours. Similar to RPE-1 cells, cilia assembly was significantly impaired upon SSNA1 KO in both 3T3 and mIMCD3 cells, indicating that SSNA1 is involved in both ciliogenesis pathways. Error bars represent the mean ± SEM. N=3. ****p < 0.0001; two-way ANOVA. **D,** Representative images of RPE-1 cells after 24-hour serum starvation. Cells were stained for the ciliogenesis suppressor CP110 (green), the mother centriole marker Ninein (magenta), the cilia marker Arl13b (red), and DNA (blue). Left: In proliferating cells, CP110 localizes to both mother and daughter centrioles, showing 2 dots at the centrosomes. Right: Upon cilia induction, CP110 is removed from the mother centriole, leaving 1 dot of signal at the centrosome. Bar, 5 µm. **E,** RPE-1 WT and SSNA1 KO cells were induced to form primary cilia by serum starvation. The ratio of CP110 removal was quantified after 12 and 24 hours of serum starvation. CP110 removal was significantly reduced in SSNA1 KO cells compared to WT, indicating that SSNA1 promotes ciliogenesis by facilitating the timely removal of CP110. Error bars represent the mean ± SEM. N=3. ****p < 0.0001; two-way ANOVA.

The assembly of cilia requires membranes to enclose the axoneme. Depending on the source of membranes, ciliogenesis can occur via the intracellular or extracellular pathway (1, 39). Intracellular pathway-dependent cells, such as RPE-1 and 3T3, rely on intracellular vesicles as the primary source of ciliary membranes. In contrast, in extracellular pathway-dependent cells, such as mIMCD3 and MDCK, the ciliary membrane is derived from the plasma membrane. Since SSNA1 promotes primary ciliogenesis in RPE-1 cells that use the intracellular pathway, we investigated whether SSNA1 is also required for the extracellular pathway. To this end, we knocked out SSNA1 in mIMCD3 cells as well as in 3T3 cells as a control mouse cell line. Similar to RPE-1 cells, cilia assembly was significantly impaired upon KO of SSNA1 in both 3T3 and mIMCD3 cells (Fig. 7*C*), from 73% in WT to 22% in KO#1 and 14% in KO#2 of 3T3 cells and from 71% in WT to 11% in KO#1 and 10% in KO#2 of mIMCD3 cells. These data indicate that SSNA1 is required for both intracellular and extracellular pathways of primary ciliogenesis.

### SSNA1 promotes ciliogenesis by facilitating the timely removal of CP110

However, regardless of the pathway used to provide membranes, the initiation of ciliogenesis also requires the removal of CP110, a key suppressor, from the centrioles. CP110 caps the distal tip of both mother and daughter centrioles in proliferating cells. As a prerequisite for cilia assembly, CP110 must be removed from the mother centriole to allow centriolar microtubules to extend and form the ciliary axoneme. To determine whether impaired ciliogenesis in SSNA1 KO cells is caused by defective CP110 removal, we stained RPE-1 cells for CP110 together with Arl13b and Ninein (Fig. 7*D*). Indeed, the efficiency of CP110 removal was significantly reduced in SSNA1 KO compared to WT cells, decreasing from 50% in WT to 17% and 18% in KO#1 and KO#2 at 12 hours after induction, and from 88% in WT to 62% and 63% in KO#1 and KO#2 at 24 hours after induction (Fig. 7*E*). These data indicate that SSNA1 promotes efficient ciliogenesis by facilitating the timely removal of the ciliogenesis suppressor CP110.

## Discussion

In summary (Fig. 8), our study challenges the previously reported roles of SSNA1 as a microtubule branching factor or nuclear antigen, showing that it is neither a MAP nor a nuclear resident protein as its name implies. Instead, using a newly developed KO-validated antibody and advanced imaging techniques, we reveal that SSNA1 is a *bona fide* centriolar protein (Fig. 8*A*). It localizes specifically to the distal lumen of both mother and daughter centrioles, as well as to the basal bodies of both primary and motile cilia. Furthermore, we uncover a hierarchical targeting network at the distal lumen of centrioles (Fig. 8*B*), where SSNA1 is arranged in a 9-fold ring pattern and interacts with both C2CD3 and LRRCC1. Although SSNA1 is not required for overall centriolar architecture, centriole biogenesis or cell division, it plays a critical role in promoting ciliogenesis (Fig. 8*C*) by facilitating CP110 removal (Fig. 8*D*).

**Fig. 8:**
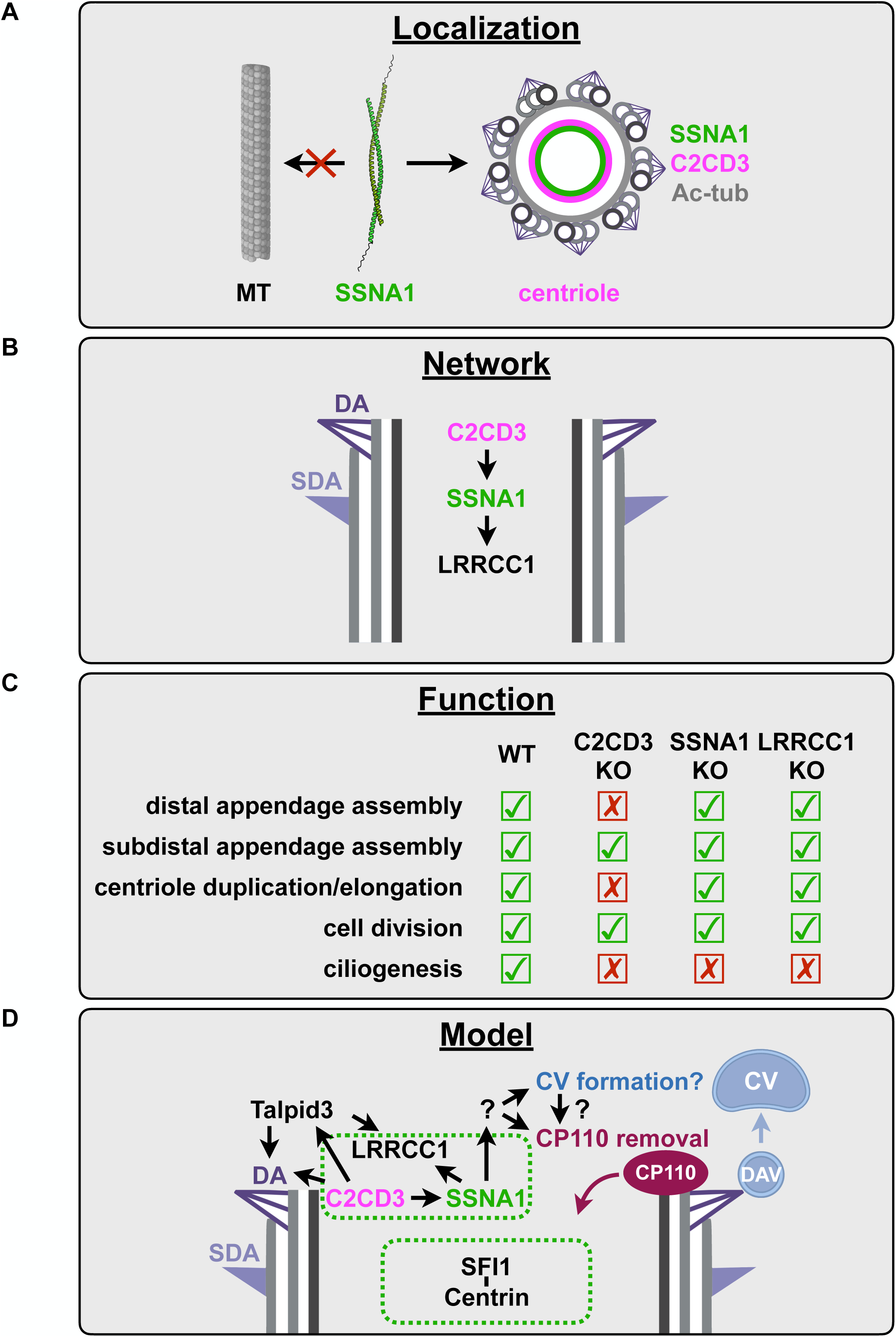
Summary of SSNA1 localization, targeting network, and physiological function, along with an integrated model to support the role of SSNA1 in distal luminal organization and ciliogenesis. **A,** Localization. SSNA1 is not a microtubule-associated protein but a genuine centriolar protein that localizes specifically to the distal lumen, where it forms a ring-like pattern and is positioned adjacent to C2CD3. **B,** Network. SSNA1 orchestrates a hierarchical assembly pathway featuring C2CD3-SSNA1-LRRCC1 at the distal lumen of centrioles. **C,** Function. SSNA1 is neither required for overall centriolar organization or duplication, nor cell division, but plays an essential role in promoting ciliogenesis. **D,** An integrated model. (1) SSNA1 organizes a distal luminal targeting network comprising the C2CD3-SSNA1-LRRCC1 module that acts separately from the SFI1-Centrin complex; (2) SSNA1 promotes ciliogenesis by facilitating the timely removal of the centriolar cap protein CP110.

SSNA1 was first identified as an autoantigen in a patient with Sjögren’s syndrome. When overexpressed in cultured cells, it appeared as numerous small puncta in the nucleus (45), hence the name of the protein. However, our results show that even minimal overexpression triggers SSNA1 self-assembly into droplets of various sizes that are distributed throughout the nucleus and cytoplasm. Upon re-examination of the original cloned cDNA sequence, we identified a sequence conflict at residue 89, reported as phenylalanine (F) (45), which differs from the NCBI reference sequence (S89) and the cDNA sequence (S89) we reverse transcribed from HEK cells. We are uncertain whether this sequence conflict simply represents a single nucleotide polymorphism (SNP) or a mutation in the original cDNA library. As we have demonstrated here, residue S89 is critical for the oligomerization state of SSNA1. Mutation of S89 to a charged residue (S89D) disrupted oligomerization, whereas substitution with a non-polar residue (S89A) produced many small droplets in the nucleus (Fig. 4*C*), closely resembling the expression pattern described for SSNA1 S89F. We believe that this sequence conflict may explain the misnomer. Nevertheless, despite its predominant centrosomal localization, excess SSNA1 can enter the nucleus due to its small size, as evidenced by the presence of nuclear droplets.

Recently, SSNA1 was further described as a microtubule remodeling and branching factor (53, 54). As shown in several reports (48, 49, 53), recombinant His-SSNA1 strongly self-associates and readily oligomerizes into high-molecular-weight filaments *in vitro*. These filaments are coated with hundreds of His residues that could weakly interact with the highly acidic C-terminal tails of tubulin. As a result, the negative electrostatic cloud on the microtubule surface may non-specifically attract SSNA1 fibrils via multivalent association with His tags. Indeed, removal of C-terminal tails of tubulin substantially weakened His-SSNA1/microtubule interaction, as reported by 49% decrease in their EDC crosslinking (53). Furthermore, by analyzing western blots of purified SSNA1 and RPE-1 cell lysates, we measured the physiological concentration of SSNA1 in RPE-1 cells to be at 2-5 nM (Fig. S11), which is four orders of magnitude lower than the concentrations used in previous *in vitro* studies (10-30 µM) (53, 54, 71). In addition to the high concentrations of recombinant proteins used, low salt (10 mM) concentration during binding reaction (54, 71) may also contribute to non-specific co-pelleting of His-SSNA1 with microtubules.

In line with this notion, we show that untagged and GST-tagged SSNA1 do not bind microtubules (Fig. 3*B*). Furthermore, removal of the N-terminal His tag or disruption of SSNA1 oligomerization by truncation or point mutation abolishes the non-specific binding of His-SSNA1 to microtubules (Figs. 3*D* and 4*D*). Indeed, the His tag is notorious for its ability to significantly enhance the binding of known MAPs to microtubules, such as EB1 (72). Moreover, the His tag can even induce artificial binding of a non-MAP to microtubules (73). For example, the recently developed His-tagged Azami-Green (AG) was designed to label microtubules *in vitro*. AG is a tetrameric fluorescent protein that does not interact with microtubules, but can be induced to artificially bind to the outer surface of microtubules after attaching a His tag to its N-terminus (73). Similar to the oligomerization of His-SSNA1, the tetramerization of His-AG (4 His tags per tetrameric AG) can therefore generate sufficient affinity for microtubules due to the multivalent interaction of His tags with tubulin. By binding and stabilizing microtubules, His-AG has been used to generate various microtubule superstructures, including branched structures, doublets, multiplets, extremely long microtubules, and asters (73). Therefore, based on multiple lines of evidence, we recommend revising the role of SSNA1 as a non-microtubule-associated centriolar protein that does not nucleate, stabilize, or branch microtubules.

When expressed in cells, SSNA1 self-assembles into droplets that in some cases can be more than 10 µm in diameter (Fig. 5*A*). We speculate that the droplets of overexpressed SSNA1 seen in primary neurons (53) may be trapped at axonal branching sites, which have the most space relative to their surrounding thin axons. These droplets may act in a dominant-negative manner, sequestering or dissociating binding partners from their native localizations (Fig. 5*A*). Thus, the actual mechanism by which SSNA1 overexpression increases axon extension and branching needs to be further investigated.

Using the newly developed SSNA1 KO cell lines and the KO-validated antibody, we investigated the subcellular localization and physiological function of SSNA1. Using various super-resolution imaging techniques, including 3D-SIM and dSTORM, in combination with ExM/TREx, we found that SSNA1 localizes to the distal lumen of centrioles and basal bodies. Due to the near molecular resolution of ExM-dSTORM, we were able to visualize the near 9-fold signal of SSNA1 arranged in a ring-like pattern. SSNA1 is positioned adjacent to but more interior than C2CD3 (Fig. 2 *C-E*) and is away from centriolar microtubules (Fig. 2*B* and *F-I*). Similarly, a gap between C2CD3 and the centriolar wall was observed in RPE-1 cells in the two recent U-ExM studies (8, 43). However, human C2CD3 and its worm homolog SAS-1 localize to the luminal side near centriolar microtubules (36, 74, 75). When expressed in human cells, worm SAS-1 becomes ectopically targeted to microtubules (74), suggesting that SAS-1 contains a microtubule-binding domain and may associate directly with centriolar microtubules. To reconcile these two seemingly contradictory observations, given that C2CD3 is a very large protein (>250 kDa), it is likely that C2CD3 adapts an extended conformation with one end interacting with centriolar microtubules and the other extending into the lumen and interacting with SSNA1 near the center. Consistent with this idea, the ring-like pattern and radial position of C2CD3 and SSNA1 from our ExM-dSTORM analysis is remarkably similar to the recently reported distal ring density RD3 by cryo-ET, with a matching diameter and location in the distal lumen of human centrioles (61). We speculate that the outer density of RD3, which is directly connected to the centriolar wall belongs to C2CD3 and that the other more inner ring-shaped density is derived from part of C2CD3 and SSNA1. In agreement, recent AlphaFold modeling of the worm SAS-1/SSNA-1 complex predicts that the C-terminus of SAS-1 interacts with SSNA-1, whereas the N-terminus of SAS-1 is located near the centriolar microtubules, away from where SSNA-1 localizes (76). Furthermore, consistent with our data on human SSNA1, in a heterologous human cell assay, worm SSNA-1 does not localize to microtubules and can only be ectopically targeted to microtubules upon co-expression with SAS-1 (76). Taken together, these results suggest that SSNA1/SSNA-1 of either species does not directly bind microtubules and that C2CD3/SAS-1 acts as an intermediate between centriolar microtubules and SSNA1/SSNA-1, echoing the cryo-ET density distribution of RD3 in human centrioles (61).

Using the droplet formation assay, we screened for SSNA1-interacting proteins at centrioles and subsequently verified their associations and targeting dependency in individual KO cells (Fig. 5). We thus uncovered a hierarchical assembly pathway consisting of a C2CD3-SSNA1-LRRCC1 cascade. Although it has long been thought that “the centriole lumen is filled with an amorphous substance at the distal end” (77), we now not only identify SSNA1 as a new component, but also establish that the distal lumen is not amorphous at all. Rather, distal luminal proteins are arranged in a specific configuration and recruited in an ordered fashion. In particular, the targeting dependency of C2CD3-SSNA1 has recently been shown to be conserved in worm (76). Furthermore, the distal lumen may accommodate two or more functional modules (61) (Fig. 8*D*), as the distal luminal complex SFI1/Centrin is not recruited to C2CD3/SSNA1/LRRCC1 droplets (Fig. 5*B*). Taken together, our results reveal an unexpected level of structural organization within the distal centriolar lumen, prompting further efforts to systematically identify additional components and analyze their spatial arrangement in this region.

Despite the conservation of its centriolar localization and upstream recruiter, SSNA1 appears to have different functions in worms and in human cells. In *C. elegans*, SSNA-1 affects centriolar stability and mitotic spindle assembly in early embryos (71, 76). However, we found that mammalian SSNA1 is dispensable for centriolar organization, biogenesis (Fig. 6), and cell division, but is required for efficient cilia assembly (Fig. 7) (78). Therefore, it will be of particular interest to further investigate its function in other organisms and/or in different biological contexts.

Distal luminal proteins such as C2CD3, SSNA1 and LRRC1 regulate ciliogenesis, although to different degrees. Our functional analyses show that ciliogenesis is most impaired in C2CD3 KO, followed by SSNA1 KO and then LRRCC1 KO (Fig. 7*B*). As described above, C2CD3 affects the recruitment of many proteins both outside and inside the distal lumen, in particular DA, which is critical for docking the mother centriole to membranes. As a result, in the absence of C2CD3, essentially no cilia could be formed. In contrast, KO of LRRCC1, which represents the most downstream component of the pathway, causes a significant but milder defect than SSNA1 KO, indicating that the more severe ciliogenesis defects induced by SSNA1 KO cannot be solely attributed to the loss of LRRCC1. Therefore, other effectors must act downstream of SSNA1 to exert full function (Fig. 8*D*). Regardless, because cilia must be present at the right time and place for proper signaling and/or motility to occur, any structural defects or kinetic delays in cilia assembly could have serious consequences *in vivo*.

During the early stage of intracellular ciliogenesis, after PCV transport and docking to the DA, the DAVs gradually fuse side-by-side to form a donut-shaped membrane structure. Subsequently, such donut-like membranes further fuse centripetally to form a large hood-like or cap-like CV that completely insulates the distal lumen (39). Simultaneously, the capping protein CP110 are removed from the tips of the centriolar microtubules, allowing the axoneme to elongate (79). Given the proximity of distal luminal proteins to the sites of CV formation and axoneme initiation, it is mechanistically tempting to speculate that distal luminal proteins may serve as sensors or effectors for the completion of a fully closed CV (Fig. 8*D*). Alternatively, to coordinate axoneme initiation with CV formation, distal luminal proteins may act as transducers, relaying signals to trigger release of the CP110 suppressor from capping microtubules. Indeed, our data indicate that CP110 removal is impaired in SSNA1 KO cells (Fig. 7*D*). Since SSNA1 is also found in the basal bodies of motile cilia, it will be interesting to determine whether and in which tissues or cell types SSNA1 may regulate assembly of motile cilia. Finally, it will be also important to investigate the tissue expression profile of SSNA1 in mammals and whether mutations in SSNA1 contribute to ciliopathies in humans (80).

## Materials and Methods

### Plasmids

All constructs were verified by Sanger sequencing and are listed in Supplementary Table 1.

#### Human SSNA1 cDNA

The full-length human SSNA1 cDNA was obtained initially through PCR amplification from a first-strand cDNA library reverse-transcribed from total mRNA isolated from HEK cells. The sequence was confirmed to match the NCBI reference sequence for SSNA1 transcript variant 1 (NM_003731.3). Coding sequences for the WT, truncated, and point mutant variants were subsequently generated by gene fragment synthesis (Twist Bioscience) or PCR-based mutagenesis.

#### Bacterial expression constructs

The bacterial expression construct for GST (pJHW1048) was modified to include a 3C cleavage site (LEVLFQ/GP) upstream of GST in the pET23a(+)-T7-GST (pJHW0082)(64) plasmid backbone.

The bacterial expression construct pET23a(+)-GM130 N74-GST (pJHW1086), which expresses the first 74 amino acids of GM130 (*i.e*., the microtubule-binding domain) fused to a C-terminal GST tag, has been described previously (64).

The bacterial expression construct for GST-SSNA1 (pJHW0896), also used for tag-cleaved SSNA1 and its variants, was generated by replacing the GFP-Nanobody coding sequence in pGEX6P1-GFP-Nanobody (Addgene #61838, pJHW0080) with the SSNA1 coding sequence. SSNA1 was positioned immediately downstream of the 3C cleavage site without introducing additional residues.

The bacterial expression construct for SSNA1-GST (pJHW1334) was generated by removing the T7 tag and inserting the SSNA1 coding sequence into pET23a(+)-T7-3C-GST (pJHW1048) upstream of 3C-GST.

To generate custom His-tagged constructs, the pET23a(+)-His-NEX-His vector (pJHW0662) was developed by replacing the T7-GST coding sequence in pET23a(+)-T7-GST (pJHW0082) with a modified multiple cloning site (MCS) containing NheI-EcoRI-XhoI (NEX) restriction sites flanked by two His tags. The pET23a(+)-His-TEV-His vector (pJHW0894) was derived by adding a TEV cleavage site (ENLYFQ/G) to pET23a(+)-His-NEX-His (pJHW0662).

The bacterial expression construct for His-SSNA1 (pJHW0895) was generated by inserting the SSNA1 coding sequence with a stop codon downstream of the TEV cleavage site into pET23a(+)-His-TEV-His (pJHW0894).

The bacterial expression construct for SSNA1-His (pJHW1338) was generated by subcloning the SSNA1 coding sequence into pET23a(+)-His-TEV-His (pJHW0894) with simultaneous removal of the His-TEV sequence.

The bacterial expression constructs for tag-cleaved SSNA1 truncated and point mutant variants (pJHW0916, pJHW1342, pJHW1349, pJHW0993, pJHW0954) were generated by subcloning the coding sequence of SSNA1 variants (residues 1-104, 1-97, or 1-90, or the S89A and S89D mutants) into the pGEX6P1-based vector with an N-terminal GST tag followed by a 3C cleavage site.

The bacterial expression constructs for His-SSNA1 oligomerization-deficient mutants (pJHW1368, pJHW1203) were generated by subcloning the coding sequence of SSNA1 variants (residues 1-90 or the S89D mutant) into the pET23a(+)-based vector with an N-terminal His tag followed by a TEV cleavage site.

#### Mammalian expression constructs

The mammalian expression vector pcDNA3.1(+)-KENNEX (pJHW0612) was custom-made by replacing the original MCS of pcDNA3.1(+) (pJHW0482) with a modified MCS containing KpnI-EcoRV-NotI-NheI-EcoRI-XhoI (KENNEX) restriction sites. Gene fragments containing coding sequences of SSNA1 WT, truncated, or point mutant variants flanked by the KpnI and XhoI restriction sites were cloned into the pcDNA3.1(+)-KENNEX vector for mammalian expression without epitope tags (pJHW0811, pJHW0874, pJHW0960, pJHW1376, pJHW0850, pJHW0939).

#### CRISPR/Cas9 constructs for human C2CD3 and human Talpid3 KO

The hCas9 expression construct (Addgene #41815) has been described previously (41). The gRNA expression constructs targeting human C2CD3 and human Talpid3 (gRNA-C2CD3_sg1_1051-1073, gRNA-Talpid3_sg1_1421-1443, gRNA-Talpid3_sg2_1465-1487) were generated by inserting annealed oligos into the gRNA cloning vector (Addgene #41824). The gRNA targeting sequences are provided in Supplementary Table 2.

#### CRISPR/Cas9 constructs for human SSNA1, mouse Ssna1 and human LRRCC1 KO

The all-in-one CRISPR vector with a GFP tag, pAll-Cas9-GFP (from the C6 RNAi Core Facility, Academia Sinica, Taiwan), was digested with BsmBI and ligated with annealed oligos. These constructs were first evaluated for the cutting efficiency of individual gRNAs. Two of the most effective gRNAs were selected to make double-cut constructs (two sets for human SSNA1, one set each for mouse Ssna1 and human LRRCC1). The final all-in-one pAll constructs (pJHW1083, pJHW1084, pJHW1432, pJHW1441) allowed simultaneous expression of Cas9, two gRNAs, puromycin N-acetyltransferase (PAC; for antibiotic selection, not applicable to RPE-1 cells) and EGFP (for cell sorting). Double cutting efficiency was also validated by a surrogate reporter assay prior to KO cell generation.

### Protein expression and purification

#### Tag-cleaved SSNA1 (WT and variants)

Plasmids were transformed into *E. coli* Rosetta2 (DE3) cells. Bacterial cultures were grown in LB medium containing appropriate antibiotics at 37 °C until the optical density (OD600) reached 0.6. Protein expression was induced with 0.25 mM IPTG, followed by overnight incubation at 16 °C. Cells were harvested by centrifugation at 6000 rpm (JLA-8.1000, Beckman) for 30 minutes at 4 °C, and the resulting pellets were resuspended in lysis buffer (50 mM potassium phosphate pH 8.0, 300 mM KCl, 5 mM β-mercaptoethanol, 2 mM EDTA). Cells were lysed using a microfluidizer, and lysates were clarified by centrifugation at 35,000 x *g* for 45 minutes at 4 °C. The supernatant was incubated with glutathione Sepharose 4B resin (Cytiva) for 2 hours at 4 °C with gentle mixing. Bound proteins were thoroughly washed and then cleaved on the resin by overnight incubation with purified 3C protease in cleavage buffer (25 mM potassium phosphate pH 7.4, 5 mM β-mercaptoethanol, 2 mM EDTA). The His-tagged 3C protease was removed using Ni-NTA beads (Qiagen). Cleaved proteins were dialyzed overnight at 4° C against 2 L of dialysis buffer (25 mM potassium phosphate pH 7.4, 0.5 mM TCEP), concentrated using a 3 kDa MWCO centrifugal device (Millipore), and quantified by Bradford assay (Bio-Rad). Size exclusion chromatography (SEC) was performed on a Superose 6 Increase 10/300 GL column (Cytiva) equilibrated in SEC buffer (25 mM HEPES pH 7.4, 150 mM NaCl, 0.5 mM TCEP or 1 mM DTT).

#### GST-SSNA1 and SSNA1-GST

Expression and purification of GST-tagged SSNA1 followed similar procedures as for tag-cleaved SSNA1, except that the bound proteins were eluted using a buffer containing 30 mM reduced glutathione instead of 3C protease cleavage.

#### His-SSNA1 (WT and variants) and SSNA1-His

Expression and purification of His-tagged SSNA1 followed similar procedures as for GST-tagged SSNA1, but with the following modifications: the lysis buffer did not contain EDTA, bound proteins were washed with a buffer containing 25 mM imidazole, and proteins were eluted with a buffer containing 250 mM imidazole instead of reduced glutathione.

### Biophysical characterization of tag-cleaved SSNA1

#### Size exclusion chromatography coupled with multi-angle light scattering (SEC-MALS)

The molecular weights of tag-cleaved SSNA1 and its oligomerization-deficient mutants (residues 1-90 and the S89D mutant) were determined by SEC-MALS. A Superdex 200 Increase 10/300 GL column (Cytiva) was connected to a DAWN HELIOS II 18-angle light scattering detector (Wyatt) and an Optilab T-rEX refractometer (Wyatt). The column was equilibrated with SEC buffer (25 mM HEPES pH 7.4, 150 mM NaCl, 0.5 mM TCEP) on an ÄKTA-UPC 900 FPLC system (Cytiva). Data were collected and processed using ASTRA v6.1.2 software (Wyatt) with Zimm plot analysis. A refractive index increment (dn/dc) value of 0.1850 ml/g, representing a standard value for proteins, was used for all samples.

#### Circular dichroism (CD) spectroscopy

Far-UV CD spectra were recorded using an AVIV model 400 circular dichroism spectrometer (Aviv Biomedical Inc). SSNA1 or variant samples (400 µl at 0.1 mg/ml) were prepared in CD buffer (25 mM potassium phosphate pH 7.4, 0.5 mM TCEP). Measurements were performed in a 1-mm pathlength quartz cuvette (Hellma) at 1 nm intervals between 260 and 190 nm. Each spectrum was recorded with a 20-second averaging time, averaged over three scans, and converted to mean residue ellipticity. Secondary structure deconvolution was performed using the BeStSel method (81).

#### Negative staining EM

A 5 µl aliquot of SSNA1 at 0.005 mg/ml in SEC buffer (25 mM HEPES pH 7.4, 150 mM NaCl, and 0.5 mM TCEP) was applied to freshly glow-discharged formvar/carbon-supported copper grids (Ted Pella). The samples were immediately stained with 5 µl of 1% (w/v) uranyl acetate. Images were captured using a Tecnai G2 Spirit TWIN transmission electron microscope at 100 keV.

### ColabFold modeling

Structural models of the SSNA1 tetramer were generated using ColabFold v1.5.2(65). High-quality models were selected based on the following cutoff criteria: predicted local distance difference test (pLDDT) score >85; and interface predicted template modeling (ipTM) score >0.6. Predicted structures were visualized and analyzed using PyMOL.

### Microtubule co-pelleting assay

The microtubule co-pelleting assay was performed as described previously (64). Briefly, microtubules were assembled from porcine brain tubulin (Cytoskeleton) at 37 °C in BRB80 buffer (80 mM PIPES pH 6.9, 1 mM MgCl2, 1 mM EGTA) supplemented with 1 mM GTP and stabilized with 20 µM taxol. Preassembled microtubules were incubated with the indicated proteins in BRB80 buffer at a final concentration of 4 µM each for 10 minutes and then pelleted through a taxol-containing cushion by centrifugation at 100,000 x *g* for 30 minutes at room temperature in a TLA100 rotor (Beckman). The resulting supernatants and pellets were analyzed by SDS-PAGE followed by Coomassie blue staining. Band intensities were quantified using the Gels function in Fiji. For TEV cleavage experiments, recombinant TEV protease was incubated with His-SSNA1 for 10 minutes at room temperature prior to the assay.

### Cell culture and transfection

#### Common cell lines and transfection

HEK (ATCC), HeLa (ATCC), and 3T3 Flp-In cells (Thermo Fisher) were cultured in Dulbecco’s modified Eagle’s medium (DMEM) supplemented with 10% cosmic calf serum and 1% Pen/Strep. hTERT RPE-1 (ATCC) and mIMCD3 Flp-In cells (Dr. Peter Jackson, Stanford) were cultured in 1:1 DMEM/F-12 medium supplemented with 10% fetal bovine serum and 1% Pen/Strep. All cell cultures were maintained at 37 °C in 5% CO2. Lipofetamine 3000 (Thermo Fisher) was used to transfect HEK, HeLa, and RPE-1 cells according to the manufacturer’s instructions. To induce cilia, RPE-1, 3T3 Flp-In or mIMCD3 Flp-In cells were arrested at G0 phase by incubation in serum-free medium for 24 hours.

### Human Nasal Epithelial Cells (HNEC)

Nasal brushes were collected from healthy human patients with the approval of the Medical Research Ethics Committee of the University Medical Center Utrecht (Tc-BIO protocols 16-586 and 22-079). Material from one donor, HNEC0268, was used in this study. HNECs were collected and differentiated as described previously (57). Briefly, HNECs were expanded on 50 μg/ml collagen IV-coated 6-well plates and cultured in basal cell (BC) expansion medium (50% BEpiCM-b, 23.5% Advanced (Ad) DMEM/F12, 20% Rspo3-Fc fusion protein conditioned medium, 10 mM HEPES, 1% GlutaMAX, 1% Pen/Strep, 2% B-27 supplement, 0.5 μg/ml hydrocortisone, 100 nM 3,3,5-triiodo-L-thyronine, 0.5 μg/ml epinephrine hydrochloride, 1.25 mM N-acetyl-L-cysteine, 5 mM nicotinamide, 1 μM A83-01, 1 μM DMH-1, 5 μM Y-27632, 500 nM SB202190, 25 ng/ml HGF, 5 ng/ml hEGF, 5 μg/ml DAPT, 25 ng/ml FGF-7, and 100 ng/ml FGF-10).

For differentiation of 2D air-liquid interface (ALI)-HNEC cultures, basal cells were dissociated using TrypLE, seeded onto 30 mg/ml PureCol-coated 24-well transwell inserts and cultured under submerged conditions in BC medium. Upon reaching confluency, the BC medium was changed to ALI-differentiation medium (50 nM A83-01, 0.5 ng/ml hEGF, 100 nM 3,3,5-triiodo-L-thyronine, 0.5 μg/ml epinephrine hydrochloride, 100 nM TTNPB (retinoic acid agonist), 0.5 μg/ml hydrocortisone, and 1% Pen/Strep in Ad DMEM/F12) supplemented with an additional 50 nM A83-01 for 3-4 days under submerged conditions. The cells were then cultured in ALI-differentiation medium supplemented with A83-01 using a minimal amount of apical culture medium to submerge the cells. After the cells reached a densely packed state, differentiation was continued in ALI-differentiation medium supplemented with 5 mM DAPT under minimal submerged conditions for approximately 15 days. The medium was refreshed twice weekly, and the apical side of the cells was washed weekly with 125 µl PBS during differentiation.

### Generation of KO cells by CRISPR/Cas9-mediated gene editing

#### C2CD3 KO and Talpid3 KO in RPE-1 cells

RPE-1 C2CD3 KO cells and Talpid3 KO cells were generated using the CRISPR/Cas9 gene editing system82 as described previously (41). Briefly, 2.5 μg of the hCas9-expressing plasmid and 2.5 µg of the specific gRNA-expressing plasmid were mixed with electroporation buffer (Lonza) according to the manufacturer’s instructions and applied to RPE-1 cells. After nucleofection, cells were serially diluted, and single colonies were then isolated and expanded.

Gene KOs were confirmed by immunofluorescence, Western blotting, and/or genomic DNA sequencing.

#### SSNA1 KO / LRRCC1 KO in RPE-1 cells and Ssna1 KO in 3T3 Flp-In / mIMCD3 Flp-In cells

The all-in-one CRISPR constructs expressing respective gRNA (Supplementary Table 2), Cas9 nuclease, and GFP were transfected either by Lipofectamine 3000 or delivered by the Neon Transfection System (Thermo Fisher) into target cells. We used 2 gRNA double cuts for SSNA1, Ssna1 and LRRCC1. Then, 48 hours post-transfection, the GFP-positive cells were subjected to fluorescence-activated cell sorting (FACS, FACSAria III, BD Biosciences) into a vial to generate a cell pool, which was used to evaluate KO efficiency, as well as into 96-well plates with one cell per well, with these latter expanded into single colonies for later screening. Briefly, genomic DNA was extracted from the cell pool or subsequently from the single expanded colonies using a genomic DNA mini kit (QIAGEN or Favorgen) according to the manufacturer’s instructions. Genotyping PCR was carried out on extracted genomic DNA and appropriate forward and reverse primers. A WT or KO PCR amplicon spanning the target site was amplified and visualized on the agarose gel to assess KO efficiency, or it was sent for Sanger sequencing. Selected KO clones were further confirmed by immunofluorescence and Western blotting analyses.

### Antibodies

#### Commercial antibodies

Information on commercial antibodies is provided in Supplementary Table 3.

#### Generation of anti-SSNA1 polyclonal and monoclonal antibodies

To generate antibodies, full-length tag-cleaved human SSNA1 proteins were used as immunogens. Polyclonal antibodies (LTK Rb pAb, YH-2005W GP pAb, and YH-2005Y GP pAb) were generated by immunizing rabbits (LTK BioLaboratories, Taiwan) and guinea pigs (Yao-Hong Biotechnology, Taiwan). The monoclonal antibody, designated clone A6, was developed by immunizing mice (LTK BioLaboratories, Taiwan) with the same immunogen. Hybridomas were subsequently generated using standard monoclonal antibody production protocols.

The A6 monoclonal antibody used in this study is an unpurified mouse ascites preparation. This monoclonal antibody has been extensively characterized and validated for specificity. It recognizes only human SSNA1, with its isotype subclass identified as mouse IgG1.

### Western blotting

RPE-1 cells were briefly washed with cold PBS and lysed on ice in TEGN buffer (10 mM Tris-HCl pH 7.4, 150 mM NaCl, 1 mM EDTA, 10% glycerol, 0.5% NP40, supplemented with a protease inhibitor cocktail). Lysates were cleared by centrifugation at 15,000 x *g* for 15 minutes at 4 °C. Total protein concentrations were measured using the Precision Red Advanced Protein Assay (Cytoskeleton). Proteins were separated by SDS-PAGE and transferred onto PVDF membranes (Millipore). After blocking with 5% milk in TBST buffer (20 mM Tris-HCl pH 7.4, 150 mM NaCl, 0.05% Tween 20) for 1 hour at room temperature, the membranes were incubated with primary antibodies for 2 hours at room temperature or overnight at 4 °C. After thorough washing, the membranes were incubated with HRP-conjugated secondary antibodies for 1 hour at room temperature. After thorough washing, the membranes were treated with home-made enhanced chemiluminescence (ECL) reagents and visualized with X-ray film.

### Imaging

#### Immunofluorescence of cell lines, epifluorescence, confocal imaging, and 3D-SIM

Cells grown on round 12 mm coverslips (No. 1 for epifluorescence and confocal imaging, No. 1.5H for 3D-SIM; Paul Marienfeld) were fixed and permeabilized in methanol for 15 minutes at -20 °C. After blocking with 1 mg/ml BSA in PBS, cells were incubated with primary antibodies for 30 minutes at 37 °C followed by Alexa Fluor-conjugated secondary antibodies for 30 minutes at 37 °C. DNA was stained with 1 µg/ml Hoechst 33342 for 5 minutes. The cells were then mounted on glass slides in home-made Mowiol 4-88 (Calbiochem) mounting solution. For 3D-SIM, the mounting solution contains 100 nm TetraSpeck fluorescent beads (1:40, Thermo Fisher, T7279) as a fiducial marker to correct for optical chromatic aberration.

While confocal imaging was employed to verify the presence of SSNA1 droplets in the nucleus, most images were taken by epifluorescence microscopy unless otherwise stated. Epifluorescence and confocal images were acquired with Plan-Apochromat 40x/1.3NA or 63x/1.4NA Oil DIC objectives on a Zeiss Axio Observer.Z1/7 system and Zen 2.6 software blue edition (Zeiss). The 3D-SIM super-resolution images were acquired using a Plan Apochromat 63x/1.4NA Oil DIC objective on a Zeiss ELYRA 7 system and ZEN software (Zeiss).

#### Immunofluorescence of airway epithelium

For immunofluorescence cell staining, the apical side of the cells was washed with 125 μl PBS at 37 °C. Cells were fixed with -20 °C methanol for 15 minutes and then washed three times for 10 minutes with PBS. Before staining, filters containing differentiated HNECs were cut out of the plastic transwell scaffold and sectioned into smaller fragments for further processing. Filter pieces were incubated with blocking buffer (3% BSA in PBS) for 1 hour. To stain the sample, filter pieces were transferred to a 0.5-ml Eppendorf tube and incubated with primary antibody in blocking buffer at 4 °C overnight, washed three times for 10 minutes with PBS, and incubated with secondary antibody in blocking buffer for 5 hours at room temperature. Then, filter pieces were washed three times for 10 minutes with PBS. Finally, filter pieces were dehydrated using a 70% EtOH wash, followed by a 100% EtOH wash, and mounted in Prolong Diamond.

#### Ten-Fold Robust Expansion (TREx) microscopy of airway epithelium

Airway epithelium was processed for expansion as described previously (58). In brief, prior to Ten-Fold Robust Expansion (TREx) microscopy, filter pieces were fixed and stained as described in the section above. After removal of the secondary antibodies, cells were post-fixed with 4% paraformaldehyde (PFA) in PBS for 10 minutes at room temperature, washed three times for 10 minutes with PBS, and incubated in 0.1 mg/ml acryloyl X-SE (AcX) with 0.01% TX100 in PBS overnight at room temperature. After filter pieces were washed twice for 15 minutes in PBS, they were placed onto a parafilm-covered glass slide with cells facing up, covered by a plastic transwell scaffold, and sealed with grease. To this gelation chamber 75 μl gelation solution (1% w/v sodium acrylate, 14.4% w/v acrylamide, 0.009% N, N’-methylenebisacrylamide, 1 x PBS, 0.1% Triton X-100, 0.15% tetramethylethylenediamine (TEMED) and ammonium persulfate (APS) in Milli-Q) was added and allowed to polymerize for 1 hour at 37 °C. The gel was removed from the plastic transwell scaffold and washed twice for 15 minutes with PBS and incubated in digestion solution (0.5% Triton X-100, 0.8 M guanidine-HCl, 7.5 U/ml Proteinase K in TAE (40 mM Tris, 20 mM acetic acid, 1 mM EDTA in Milli-Q)) for 4 hours at 37 °C. After three washes in PBS of 15 minutes each, the sample was incubated in 20 μg/ml ATTO-NHS-594 (Atto-Tec, AD 594-31) in PBS for 1.5 hours at room temperature. Then, 5 μg/ml of DAPI was added during the last 15 minutes of incubation. To expand the sample, gels were transferred to a 15-cm petri dish and incubated in Milli-Q at room temperature. The Milli-Q was refreshed after 30 minutes, and the sample was expanded overnight at room temperature. Prior to imaging, the cells were trimmed and mounted onto a plasma-cleaned, poly-L-lysine-treated cover glass.

Data were acquired on a Leica TCS SP8 STED3X microscope with a 405-nm DMOD Flexible and pulsed (80 MHz) white-light lasers, as well as PMT and HyD detectors. Immunofluorescence images were acquired using a HC PL APO 93x/1.3 GLYC motCORR STED objective, whereas expansion samples were acquired using a HC PL APO 86x/1.20W motCORR STED objective.

#### ExM-dSTROM imaging

Ex-dSTORM imaging was conducted to visualize centriolar ultrastructure, as described previously (36). Cells fixed with ice-cold methanol on coverslips were treated with an infusion solution (1.4% formaldehyde, 2% acrylamide in PBS) at 37 °C for 5 hours, followed by PBS washes. A gelation solution (19% sodium acrylate, 10% acrylamide, 0.1% bis-acrylamide, 0.5% TEMED, 0.5% APS in PBS) was applied on ice for 1 minute, then incubated at 37 °C for 1 hour. Hydrogels were treated with a denaturation buffer (200 mM NaCl, 200 mM SDS, 50 mM Tris-HCl pH 9.0), boiled at 95 °C for 90 minutes, and expanded in ddH2O before immunostaining. Primary and secondary antibodies were incubated at 37 °C for 3 hours with gentle shaking. A marker protein (ATP synthase, ab109867, Abcam), labeled with Alexa Fluor 488 (1:100 dilution), was used for *in situ* drift correction. Gels were fully expanded in ddH2O at room temperature.

For dSTORM imaging, the specimen-gel composites were re-embedded in a solution (10% acrylamide, 0.15% bis-acrylamide, 0.05% TEMED, 0.05% APS) and incubated twice for 25 minutes each with shaking. Polymerization occurred in a nitrogen-filled chamber at 37 °C for 1.5 hours, followed by washing. Samples were placed in a custom-made holder and immersed in imaging buffer (Tris-HCl, NaCl, 10 mM mercaptoethylamine, 10% glucose, 0.5 mg/ml glucose oxidase, 40 μg/ml catalase).

dSTORM imaging was performed using a custom-built system based on an inverted microscope (Nikon Eclipse Ti2-E) with controlled light sources. Epi-illumination was used for dSTORM with beams from 637 nm, 561 nm, 488 nm, and 405 nm lasers and focused onto a 100x oil-immersion objective (1.49 NA). The 637 nm and 561 nm lasers were operated at high intensity (2-5 kW/cm²) to quench fluorescence, whereas the 405 nm laser converted fluorophores to a fluorescent state. The 488 nm laser was activated every 800 frames for drift correction.

Fluorescent emissions were filtered using a quad-band filter (ZET405/488/561/640 mv2, Chroma) and detected by an EMCCD camera (iXon Life 888, Andor, 83.5 nm pixel size). Each image was composed of 15,000 to 30,000 frames captured at 50 frames per second. Marker signals were used for drift correction and later removed using custom algorithms (Labview, Matlab, ImageJ) before localizing single-molecule peaks with the MetaMorph Superresolution Module (Molecular Devices).

## Data availability

All data that support the findings of this study are available within the article and its supplementary materials.

## Acknowledgments

We thank Dr. Sam Li (UCSF) and Dr. Joachim Seemann (UT Southwestern) for critical reading of the manuscript and insightful comments. We thank Dr. Naoko Mizuno (NIH) for generously sharing commercial anti-SSNA1 antibodies and mouse Ssna1 plasmids to initiate the project. We thank Dr. Pierre Gönczy for communicating results prior to publication. We thank Chuang-Kai Chueh, Jhih-Jie Tsai, the Imaging Core Facilities (Institute of Molecular Biology and Institute of Biomedical Sciences, Academia Sinica), the Biophysics Core and the RNA Technology Platform and Gene Manipulation Core (Institute of Molecular Biology, Academia Sinica) for technical support. J.H.W. is supported by the Academia Sinica Career Development Award (AS-CDA-109-L02), the Ministry of Science and Technology 2030 Cross-Generation Excellent Young Scholars Award (MOST 111-2628-B-001-013-MY3), a National Science and Technology Council Grant (NSTC 113-2113-M-002-020-), and intramural funding from the Institute of Molecular Biology, Academia Sinica, Taiwan. T.T.Y. is supported by National Science and Technology Council Grants (NSTC 112-2628-E-002-028-and 113-2628-E-002-014-) and funding from the Center for Advanced Computing and Imaging in Biomedicine (NTU-113L900703) of The Featured Areas Research Center Program within the framework of the Higher Education Sprout Project of the Ministry of Education (MOE) in Taiwan. A.A. and J.M.B. are supported by the Netherlands Organization for Scientific Research (NWO) Gravitation programme IMAGINE! (project number 24.005.009).

## Contributions

J.H.W. conceived and designed the experiments, analyzed the data, and wrote the manuscript.

Y.C.H. and X.J.C. performed most of the experiments not mentioned below. T.B.C. and T.T.Y. performed ExM-dSTORM imaging on RPE-1 cells. E.G. and A.A. performed TREx-confocal imaging on multiciliated HNECs. W.J.C. characterized SSNA1 antibodies and generated SSNA1 KO and LRRCC1 KO cells. W.B.H. and T.K.T. generated C2CD3 KO and Talpid3 KO cells.

J.M.B provided HNECs, essential expertise and reagents for multicilia induction.

---

**Fig. S1:**
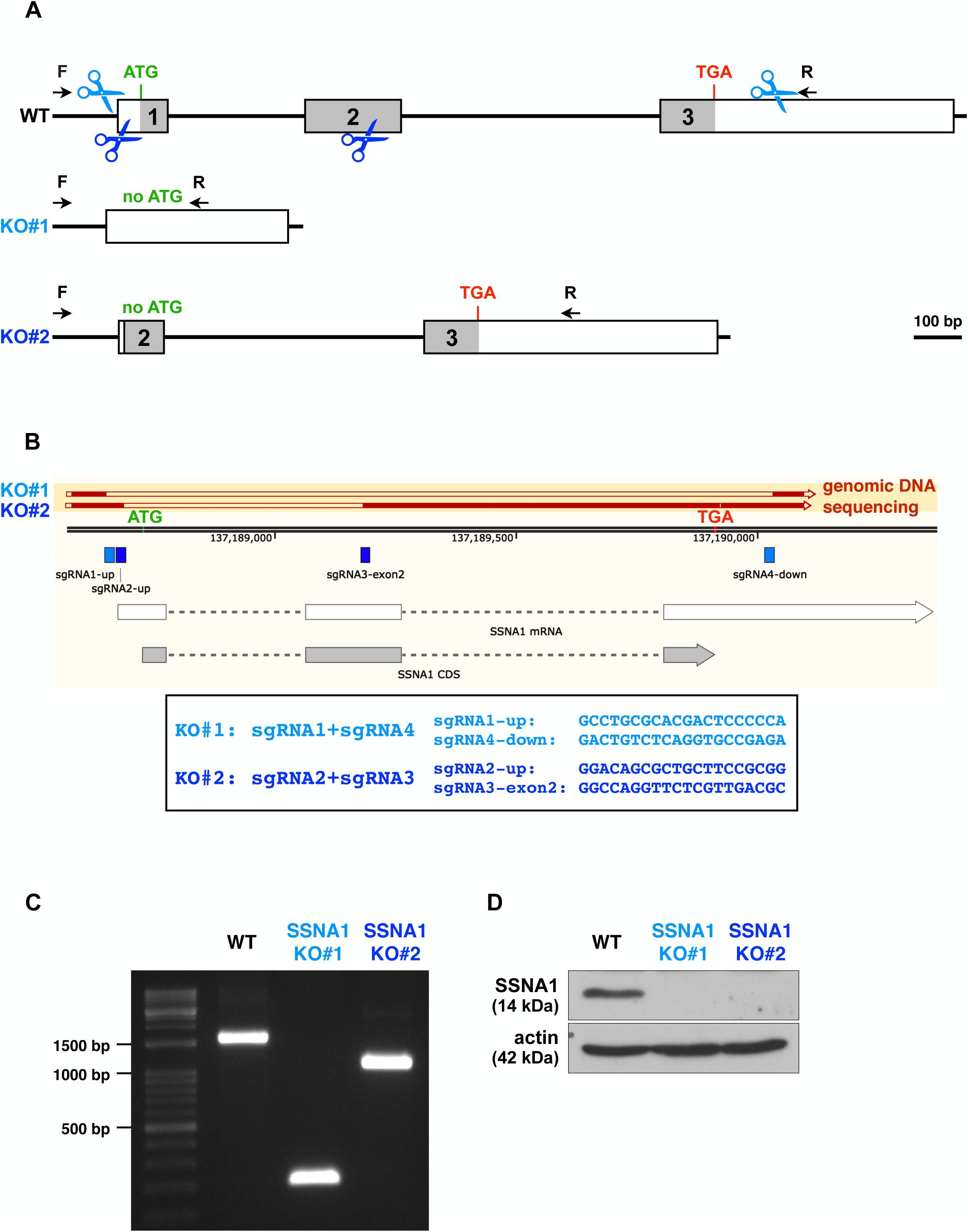
SSNA1 KO strategy and validation. **A,** Schematic diagram showing SSNA1 gene organization and the KO strategy. To ensure reliability, two different KO designs using a double-cut strategy were implemented, resulting in two independent clones, KO#1 and KO#2. The first cut was placed before the ATG start codon, resulting in no protein translation. **B,** Schematic diagram of the SSNA1 gene locus, including start (ATG in green) and stop (TGA in red) codons. Sanger sequencing results of the genomic DNA extracted from SSNA1 KO#1 and SSNA1 KO#2 cells have been aligned into the map, showing that respective genomic fragments were cut out. sgRNA sequences are also shown on the map to indicate cutting positions. **C,** Agarose gel showing genotyping PCR products from RPE-1 WT, SSNA1 KO#1 and SSNA1 KO#2 cells using the forward and reverse primers indicated in **A**. **D,** Western blots showing an absence of SSNA1 protein in both SSNA1 KO#1 and SSNA1 KO#2 cells. SSNA1 was probed with PTG rabbit polyclonal antibodies. Actin served as a loading control.

**Fig. S2:**
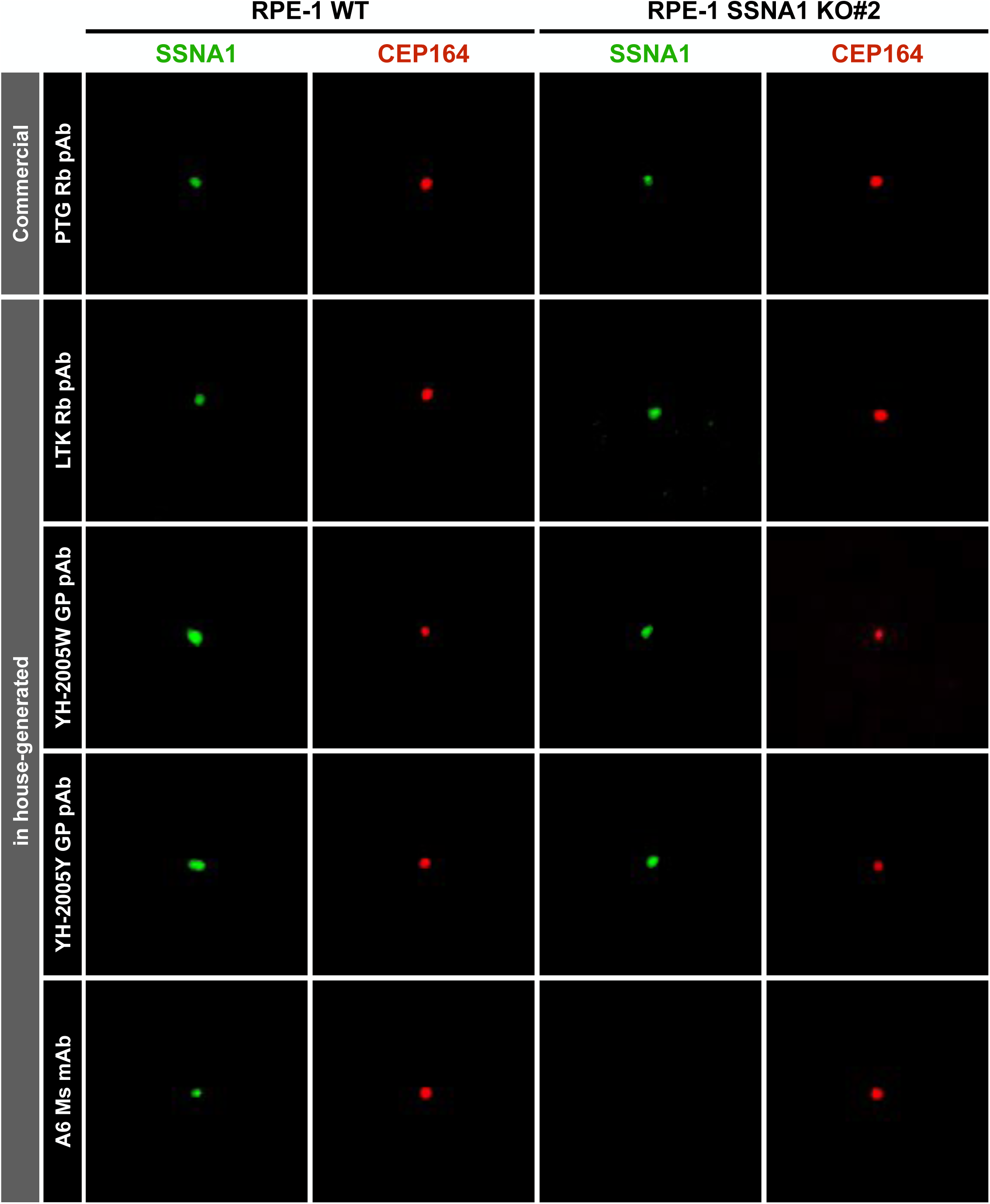
Characterization of SSNA1 polyclonal and monoclonal antibodies. Commercially available rabbit polyclonal antibodies (PTG) confirmed the absence of the expected 14 kDa SSNA1 band in KO cells by Western blotting (Fig. S1*D*), but still stained strongly at centrosomes by immunofluorescence, indicating that the PTG antibodies cross-reacted with other centrosomal proteins. To obtain SSNA1-specific antibodies, several polyclonal antibodies against SSNA1 were raised in rabbits (Rb) or guinea pigs (GP) using tag-free full-length human SSNA1 as immunogen. All polyclonal antibodies showed cross-reactivity and gave strong centrosomal signals in SSNA1 KO cells by immunofluorescence, similar to the commercial PTG polyclonal antibodies. To this end, a mouse monoclonal antibody clone A6 was developed with hybridoma technology. A6 specifically recognized SSNA1 by immunofluorescence, as the signal completely disappeared in both SSNA1 KO cell lines. CEP164 served as a positive centrosomal marker.

**Fig. S3:**
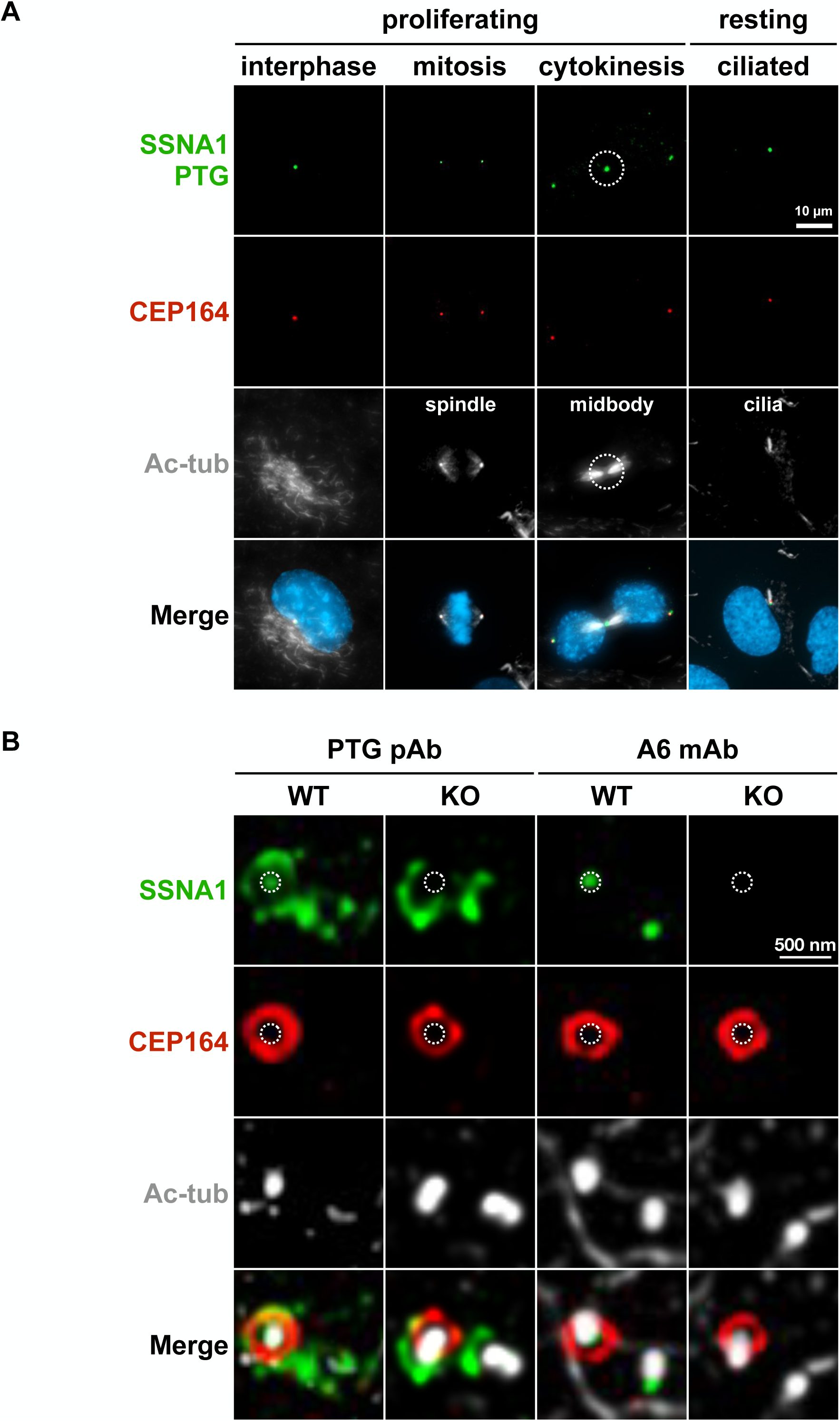
Cross-reactivity of commercial SSNA1 rabbit polyclonal antibodies. **A,** Localization of SSNA1 in proliferating and resting RPE-1 cells. Exponentially growing and starvation-induced ciliated RPE-1 cells were stained with antibodies against SSNA1 (PTG), the centrosomal marker CEP164, and acetylated tubulin (Ac-tub), which labels stable microtubules in interphase, spindle microtubules in mitosis, the midbody in cytokinesis, and the axoneme in ciliated cells. In addition to a centrosomal localization, PTG rabbit polyclonal antibodies non-specifically stained midbodies (circled) in cytokinesis (vs. Fig. 1, A6 mouse monoclonal antibody), indicating that the previously reported midbody localization was caused by antibody cross-reactivity. Bar, 10 µm. **B,** Representative 3D-SIM images of the suborganellar localization of SSNA1 at centrosomes. Exponentially growing RPE-1 cells were stained with antibodies against SSNA1 (PTG), the mother centriole/DA marker CEP164, and the centriolar microtubule marker acetylated tubulin (Ac-tub). 3D-SIM revealed multiple signals labeled by PTG antibodies at centrosomes in WT cells, with one spot located within the CEP164 ring (circled) and others appearing as satellite-like puncta around the two centrioles. In SSNA1 KO cells, the signal within the CEP164 ring (circled) was absent, suggesting that this may represent the true localization of SSNA1, while the surrounding puncta remined, which were attributable to antibody cross-reactivity. Bar, 500 nm.

**Fig. S4:**
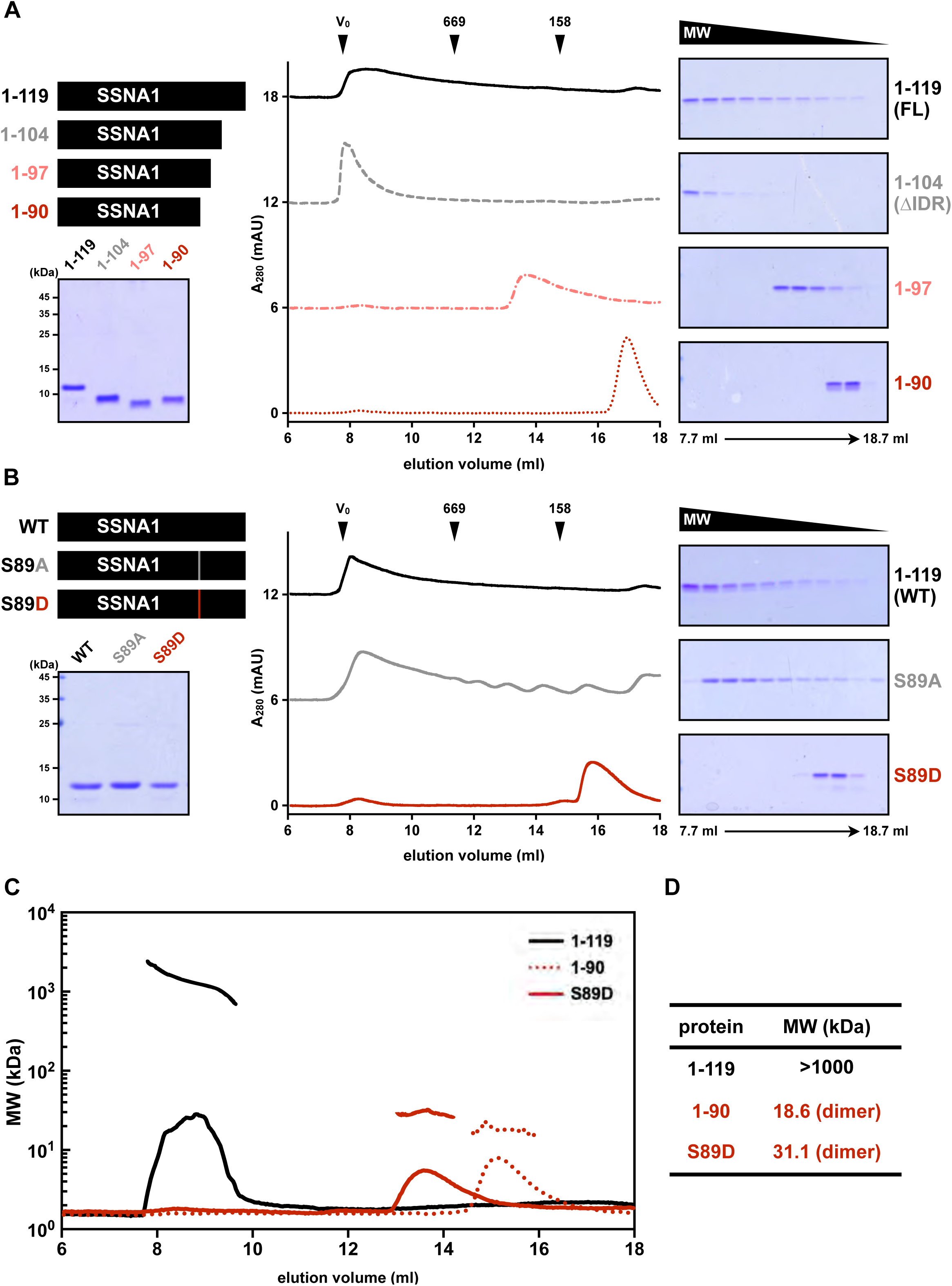
Size exclusion chromatography (SEC) purification of tag-cleaved SSNA1 and the molecular weight determination of two oligomerization-deficient mutants by SEC-MALS. Recombinant GST-SSNA1 was bacterially expressed and affinity purified, followed by 3C protease cleavage and SEC purification. The resulting tag-cleaved SSNA1 retained only two additional amino acids (Gly-Pro) at the N-terminus, closely resembling native SSNA1 as found *in vivo*. **A,** Left: schematic diagram of the SSNA1 truncated constructs and purified proteins on the Coomassie blue-stained gel. Right: SEC purification profiles of the indicated proteins. **B,** Left: schematic diagram of the SSNA1 S89 point mutation constructs and purified proteins on the Coomassie blue-stained gel. Right: SEC purification profiles of the indicated proteins. **C,** SEC-MALS analysis of two oligomerization-deficient mutant proteins, 1-90 and S89D. **D,** Molecular weight determination of 1-90 and S89D, showing that both oligomerization-deficient mutants formed dimers.

**Fig. S5:**
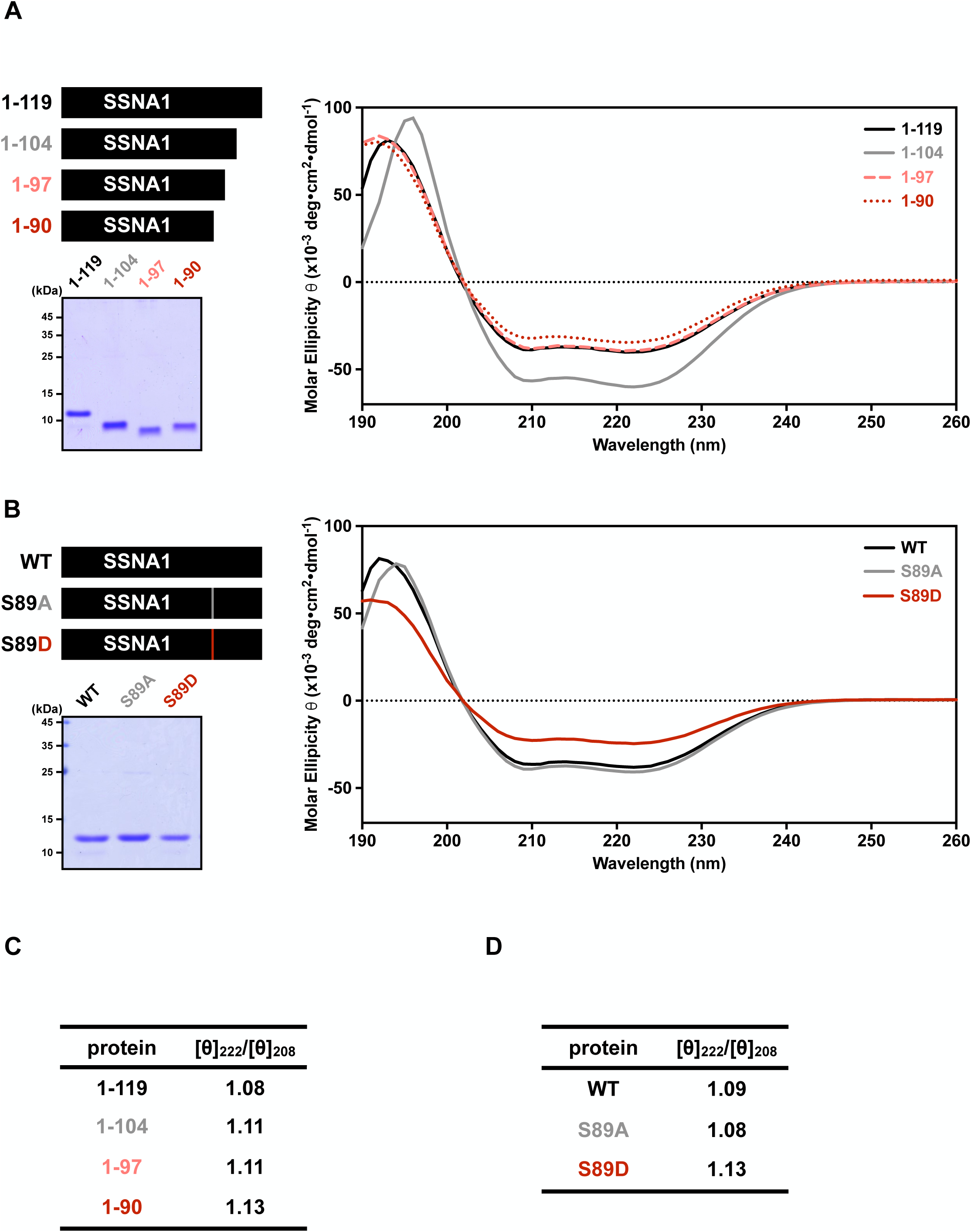
CD analysis of tag-cleaved SSNA1. All tag-cleaved SSNA1 variants showed a typical CD spectrum characteristic of α-helical profiles, with a positive band around 190 nm and two negative minima at 208 nm and 222 nm. The molar ellipticity ratio at 222 nm to 208 nm (222/208) was greater than one for all proteins, indicating that the coiled-coil structure formed in all truncated and S89 mutant variants. **A,** Left: schematic diagram of the SSNA1 truncated constructs and purified proteins on the Coomassie blue-stained gel. Right: CD spectrum of the indicated proteins. **B,** Left: schematic diagram of the SSNA1 S89 point mutation constructs and purified proteins on the Coomassie blue-stained gel. Right: CD spectrum of the indicated proteins. **C-D,** The molar ellipticity ratio [θ]222/[θ]208 of indicated proteins. The ratio was greater than one, indicating that their α-helices are in a coiled-coil conformation instead of single helix, supporting the dimeric state of the oligomerization-deficient mutants 1-90 and S89D.

**Fig. S6:**
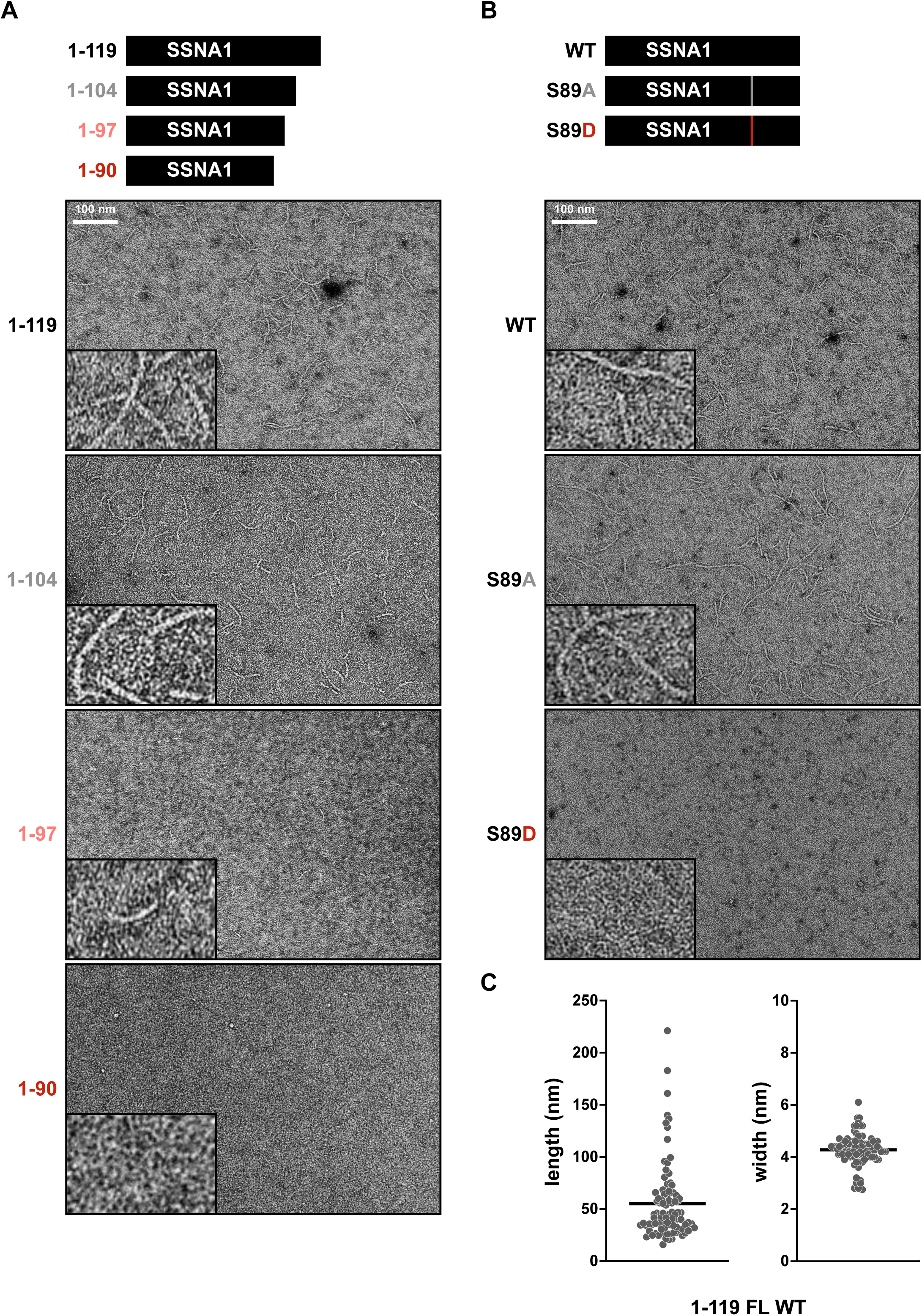
Negative-staining EM analysis of tag-cleaved SSNA1. Tag-cleaved SSNA1 was negatively stained and examined under TEM to investigate its ability to self-assemble into filaments *in vitro*. **A,** Schematic diagram of the SSNA1 truncated constructs and their representative negative-staining EM images. Bar, 100 nm. Representative images at 4x magnification are shown in the insets. **B,** Schematic diagram of the SSNA1 S89 point mutant constructs and their representative negative-staining EM images. Bar, 100 nm. Representative images at 4x magnification are shown in the insets. **C,** Quantification of length and width of WT SSNA1 filaments.

**Fig. S7:**
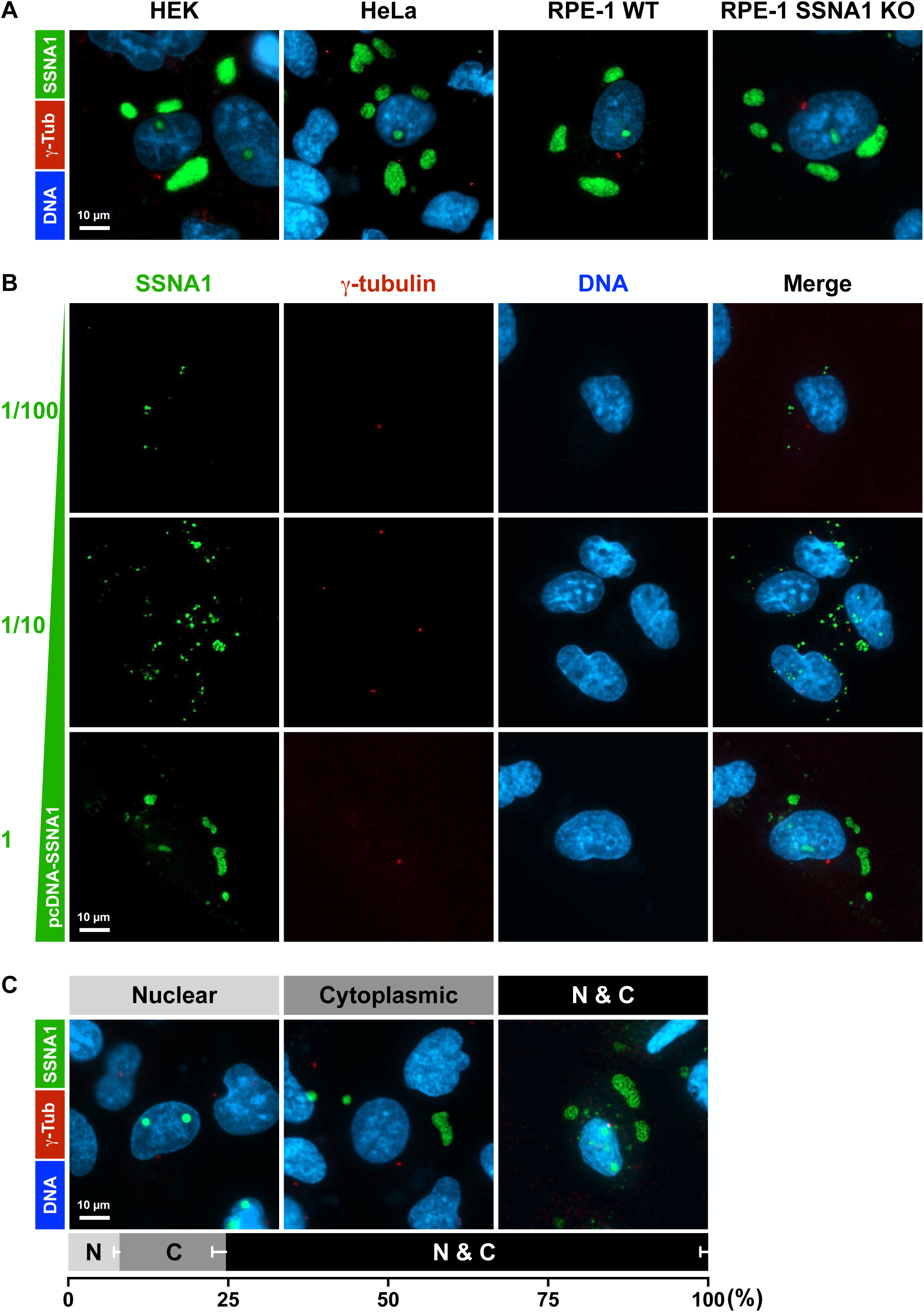
Effects of SSNA1 overexpression in cells. **A,** A mammalian expression construct containing CMV promoter-driven untagged full-length human SSNA1 was transfected into four cell lines, including HEK, HeLa, RPE-1 WT, and RPE-1 SSNA1 KO cells. Transfected cells were stained for SSNA1 (A6), the centrosomal marker γ-tubulin, and DNA. Overexpressed SSNA1 formed droplet-like structures in cells. Large droplets, when present in the nucleus, were often excluded from DNA-rich regions. Bar, 10 µm. **B,** Droplet size varied considerably, depending on SSNA1 expression levels. Although at low expression levels SSNA1 occasionally appeared as small dots near the centrosomal marker γ-tubulin, the majority of exogenously overexpressed SSNA1 did not localize to centrosomes. Bar, 10 µm. **C,** Top: Self-assembled SSNA1 droplets formed in both the nucleus and cytoplasm. Bottom: Quantification of droplet localization. 8.0% of SSNA1-overexpressing cells contained droplets only in the nucleus, 16.6% only in the cytoplasm, and 75.4% had droplets in both compartments. Bar, 10 µm.

**Fig. S8:**
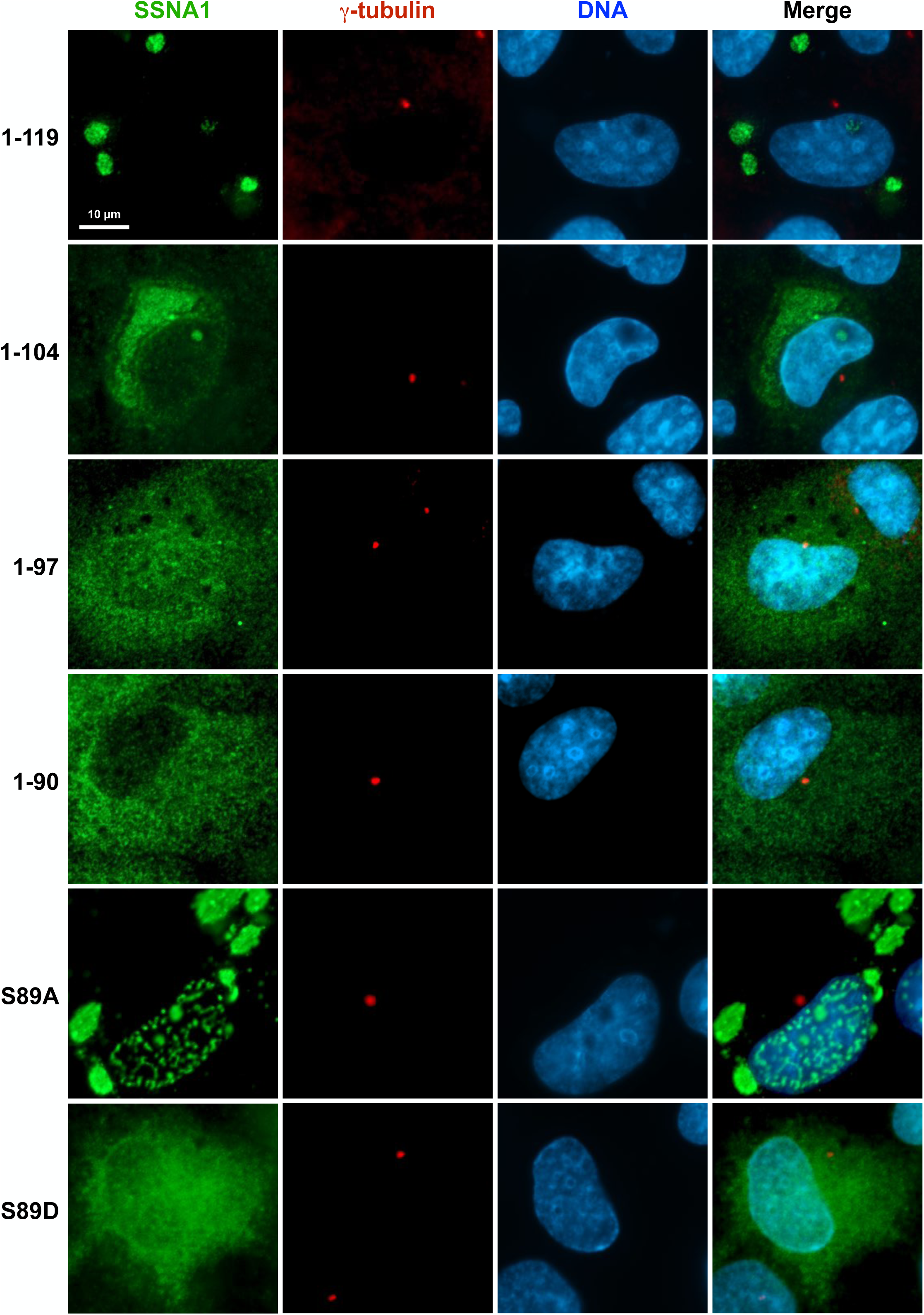
Effects of overexpressing SSNA1 variants in cells. Mammalian expression constructs containing CMV promoter-driven untagged human SSNA1, including full-length WT (1–119), truncated variants (1-104, 1-97, and 1-90), and S89 point mutants (S89A and S89D), were transfected into RPE-1 SSNA1 KO cells. Transfected cells were stained for SSNA1 (PTG), the centrosomal marker γ-tubulin, and DNA. Note that the PTG rabbit polyclonal antibodies were used in this set of experiments. Since the epitope of the A6 mouse monoclonal antibody is located within the C-terminal 15 residues, the truncated variants could not be detected by A6 antibody. Bar, 10 µm. Overexpressed SSNA1 1-119 and 1-104 formed droplets, whereas the 1-97 and 1-90 truncated variants were dispersed throughout the cells. S89A formed droplets, whereas S89D was dispersed throughout the cells. The ability of SSNA1 to form droplets in cells was correlated with its ability to form filaments *in vitro* (Fig. S6). Two oligomerization mutants, 1-90 and S89D, were confirmed as failing to form droplets.

**Fig. S9:**
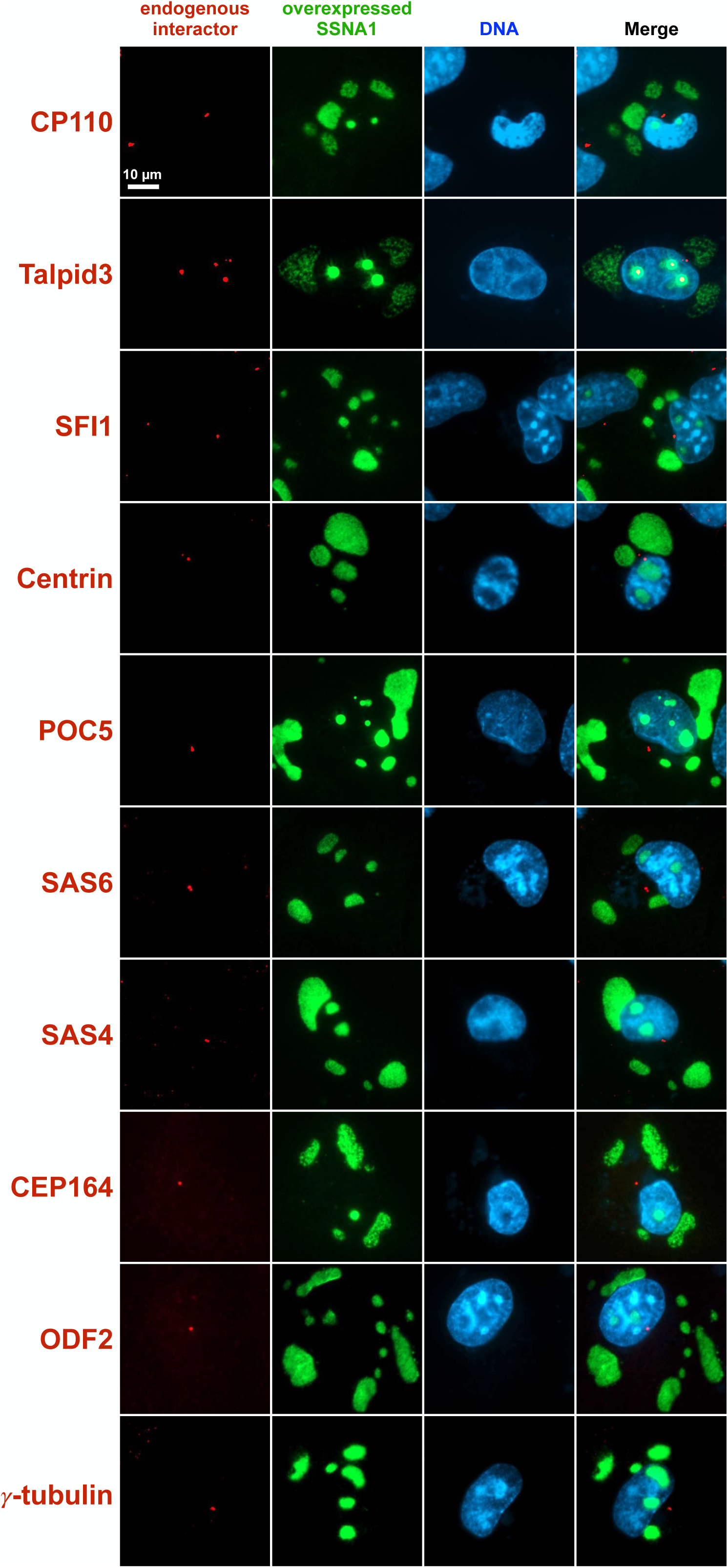
SSNA1 droplet formation and colocalization assay. Untagged full-length SSNA1 was overexpressed in SSNA1 KO RPE-1 cells. Transfected cells were stained with anti-SSNA1 A6 antibody to locate the droplets and with the antibodies against candidate proteins to test for potential interactions. As a control, most centrosomal markers representing different regions of centrosomes, including the axoneme cap (CP110), distal lumen (SFI1, Centrin), central lumen (Centrin, POC5), proximal end/procentriole (SAS6, SAS4), DA (CEP164), SDA (ODF2), and PCM (γ-tubulin), were not incorporated into SSNA1-positive droplets, indicating no interaction with SSNA1. For Talpid3, only a small percentage of cells showed a low degree of colocalization, which may have been indirectly pulled into the droplets by its interacting partner C2CD3.

**Fig. S10:**
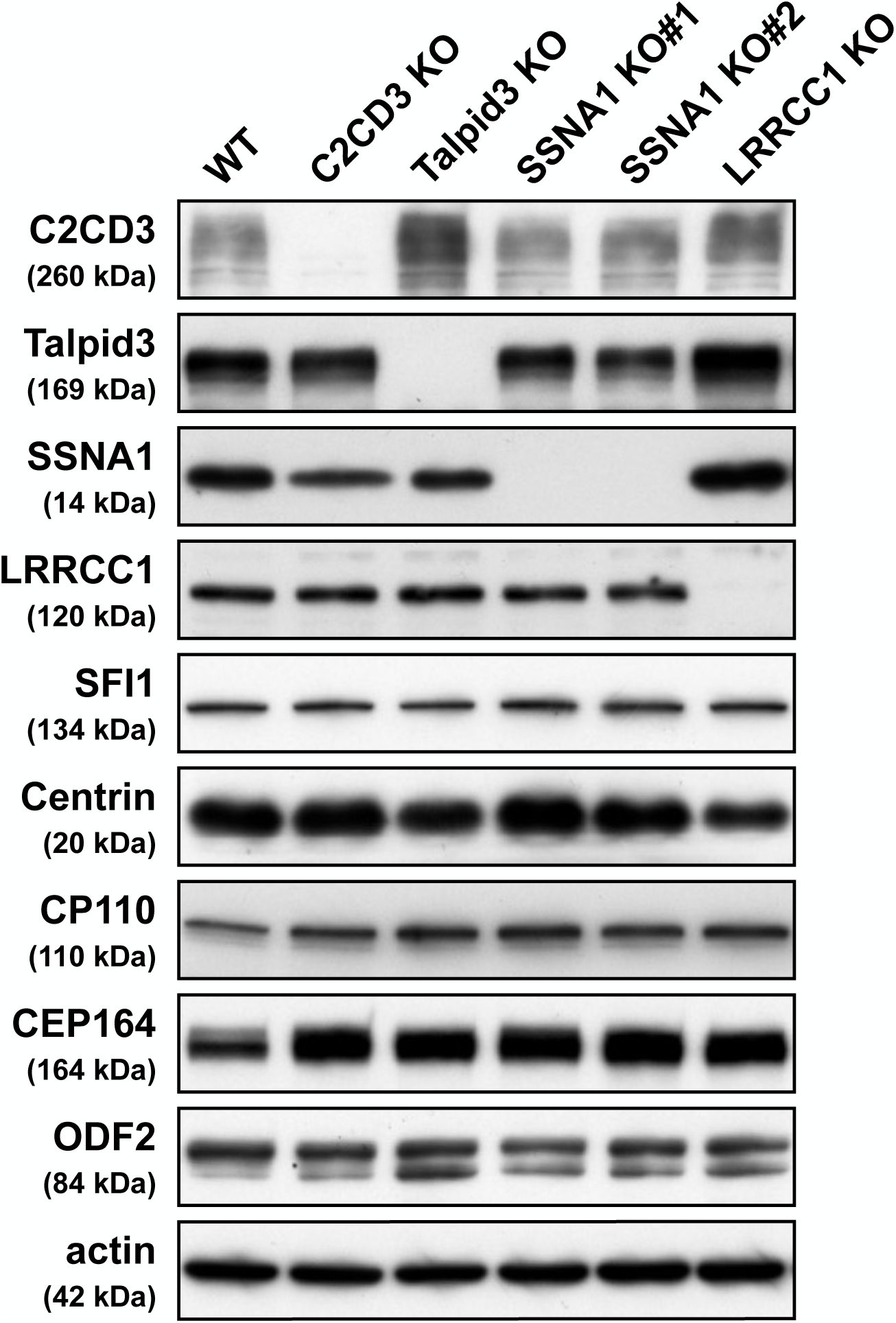
Levels of indicated proteins in RPE-1 WT and individual KO cells. Western blot analysis of total cell lysates from RPE-1 WT, C2CD3 KO, Talpid3 KO, SSNA1 KO#1, SSNA1 KO#2 and LRRCC1 KO cells, blotted using anti-C2CD3, Talpid3, SSNA1, LRRCC1, SFI1, Centrin, CP110, CEP164, ODF2, and actin antibodies.

**Fig. S11:**
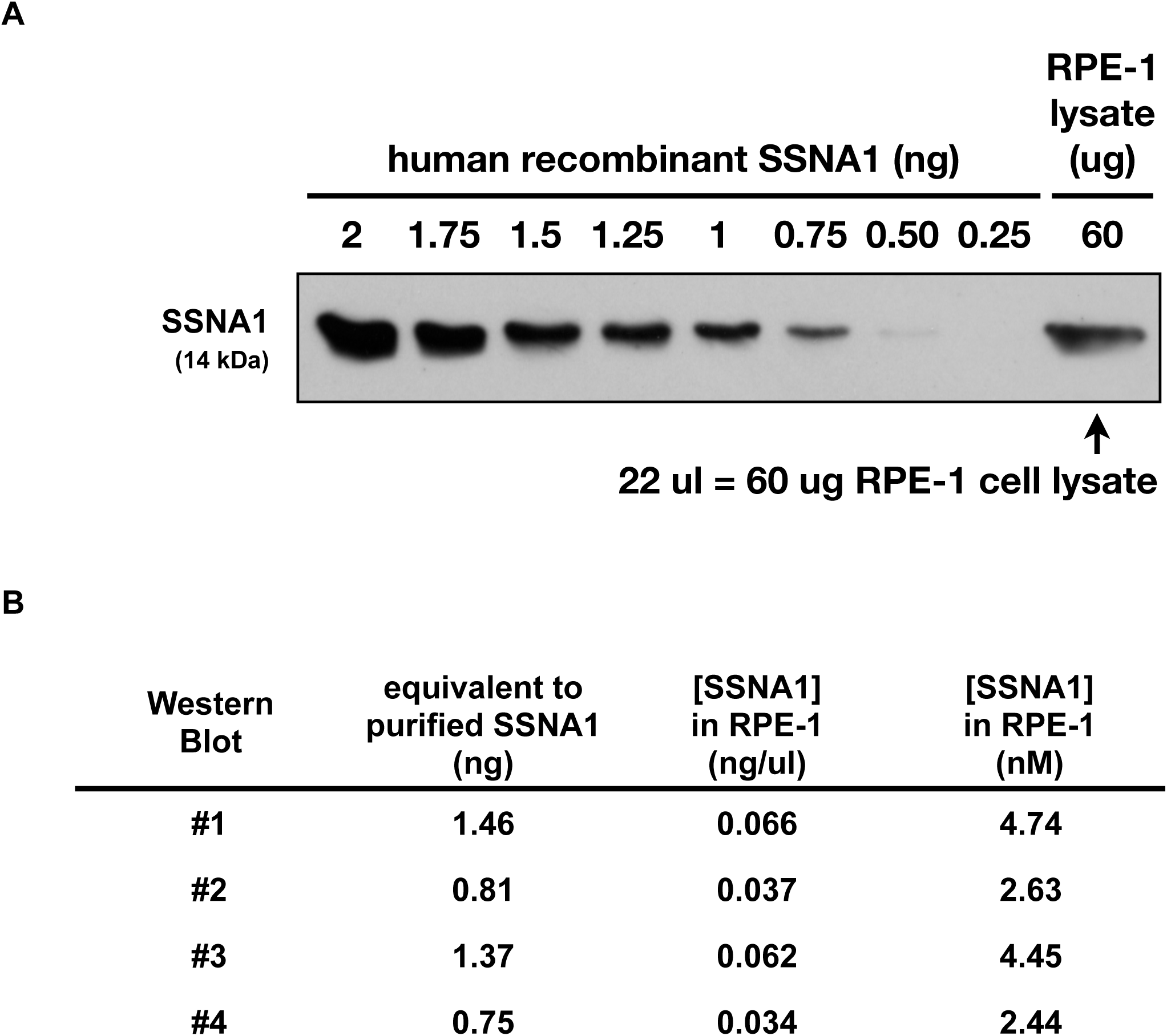
Determination of the cellular concentration of SSNA1 in RPE-1 cells. **A,** A representative image of a Western blot showing different amounts (0.25-2 ng) of recombinant tag-cleaved human SSNA1 together with 60 µg of total lysates from RPE-1 cells. **B,** Quantitation of four Western blot analyses, estimating the cellular concentration of SSNA1 in RPE-1 cells to be at 2-5 nM. N=4.

**Supplementary Table 1:** Plasmids.

| Plasmid # | Name | Ref / Addgene # |
| --- | --- | --- |
| **Bacterial expression** |  |  |
| pJHW0082 | pET23a(+)-T7-GST | Wei et al. |
| pJHW1048 | pET23a(+)-T7-3C-GST |  |
| pJHW1086 | pET23a(+)-GM130 N74-GST | Wei et al. |
| pJHW0080 | pGEX6P1-GFP-Nanobody | Addgene # 61838 |
| pJHW0896 | pGEX6P1-3C-SSNA1_WT |  |
| pJHW1334 | pET23a(+)-SSNA1-3C-GST |  |
| pJHW0662 | pET23a(+)-His-NEX-His |  |
| pJHW0894 | pET23a(+)-His-TEV-His |  |
| pJHW0895 | pET23a(+)-His-TEV-SSNA1_WT |  |
| pJHW1338 | pET23a(+)-SSNA1-His |  |
| pJHW0916 | pGEX6P1-3C-SSNA1_1-104 |  |
| pJHW1342 | pGEX6P1-3C-SSNA1_1-97 |  |
| pJHW1349 | pGEX6P1-3C-SSNA1_1-90 |  |
| pJHW0993 | pGEX6P1-3C-SSNA1_S89A |  |
| pJHW0954 | pGEX6P1-3C-SSNA1_S89D |  |
| pJHW1368 | pET23a(+)-His-TEV-SSNA1_1-90 |  |
| pJHW1203 | pET23a(+)-His-TEV-SSNA1_S89D |  |
| pJHW0438 | MBP-3Csite-3C-His |  |
| pJHW0443 | MBP-TEVsite-His-TEV | Addgene # 8827 |
| **Mammalian expression** |  |  |
| pJHW0482 | pcDNA3.1(+) |  |
| pJHW0612 | pcDNA3.1(+)-KENNEX |  |
| pJHW0811 | pcDNA3.1-KX-SSNA1_WT |  |
| pJHW0874 | pcDNA3.1-KX-SSNA1_1-104 |  |
| pJHW0960 | pcDNA3.1-KX-SSNA1_1-97 |  |
| pJHW1376 | pcDNA3.1-KX-SSNA1_1-90 |  |
| pJHW0850 | pcDNA3.1-KX-SSNA1_S89A |  |
| pJHW0939 | pcDNA3.1-KX-SSNA1_S89D |  |
| **CRISPR** |  |  |
| n/a | hCas9 | Addgene #41815 |
| n/a | gRNA_Cloning Vector | Addgene #41824 |
| n/a | gRNA-C2CD3_sg1_1051-1073 |  |
| n/a | gRNA-Talpid3_sg1_1421-1443 |  |
| n/a | gRNA-Talpid3_sg2_1465-1487 |  |
| pJHW1083 | pAll-Cas9-P2A-EGFP-F2A-Puro_(LoxP)-SSNA1_sg1+4 |  |
| pJHW1084 | pAll-Cas9-P2A-EGFP-F2A-Puro_(LoxP)-SSNA1_sg2+3 |  |
| pJHW1432 | pAll-Cas9-P2A-EGFP-F2A-Puro_(LoxP)-Mus Ssna1_sg2+4 |  |
| pJHW1441 | pAll-Cas9-P2A-EGFP-F2A-Puro_(LoxP)-LRRCC1_sg1+4 |  |

**Supplementary Table 2:**
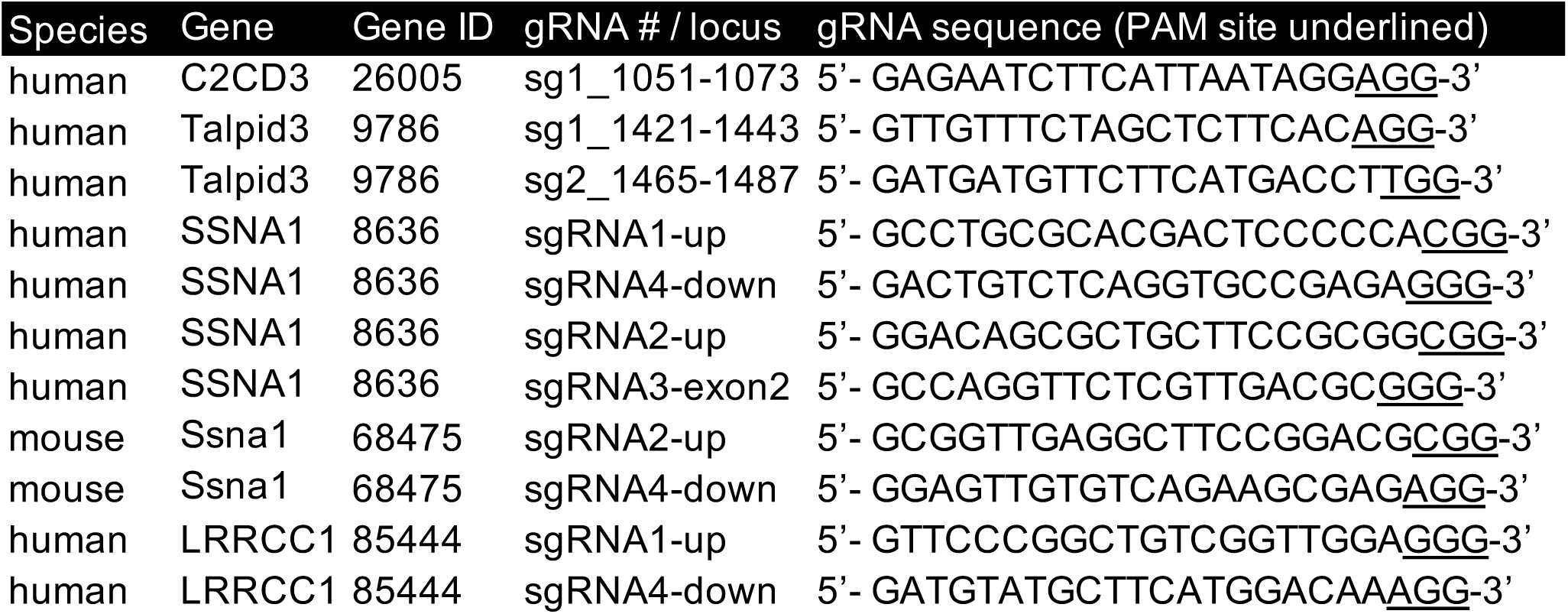
Guide RNA sequences.

**Supplementary Table 3:** Primary Antibodies.

| Primary Antibody | Supplier | Catalog # | Host | Dilution | RRID |
| --- | --- | --- | --- | --- | --- |
| WB |  |  |  |  |  |
| C2CD3 | MPS | HPA038552 | Rb | 1:300 | AB_10669542 |
| Talpid3 | PTG | 24421-1-AP | Rb | 1:1500 | AB_2879539 |
| SSNA1 | PTG | 11797-1-AP | Rb | 1:500 | AB_2195497 |
| LRRCC1 | MPS | HPA075880 | Rb | 1:750 | n/a |
| SFI1 | PTG | 13550-1-AP | Rb | 1:1000 | AB_10596938 |
| Centrin | PTG | 15877-1-AP | Rb | 1:1000 | AB_2077383 |
| CP110 | PTG | 12780-1-AP | Rb | 1:500 | AB_10638480 |
| CEP164 | PTG | 22227-1-AP | Rb | 1:1000 | AB_2651175 |
| ODF2 | MPS | HPA001874 | Rb | 1:1000 | AB_1079522 |
| actin | MPS | A5441 | Ms IgG1 | 1:100000 | AB_476744 |
| IF |  |  |  |  |  |
| SSNA1_PTG | PTG | 11797-1-AP | Rb | 1:500 | AB_2195497 |
| SSNA1_LTK | LTK | LTK-01 | Rb | 1:1000 | n/a |
| SSNA1_2005W | YH | YH-01 | GP | 1:2000 | n/a |
| SSNA1_2005Y | YH | YH-02 | GP | 1:2000 | n/a |
| SSNA1_A6 | LTK | WT#A6 | Ms IgG1 | 1:5000 | n/a |
| C2CD3 | R&D | AF7348 | Shp | 1:200 | AB_10997138 |
| Talpid3 | PTG | 24421-1-AP | Rb | 1:1500 | AB_2879539 |
| LRRCC1 | MPS | HPA012893 | Rb | 1:100 | AB_1852434 |
| LRRCC1 | MPS | HPA075880 | Rb | 1:100 | n/a |
| SFI1 | PTG | 13550-1-AP | Rb | 1:200 | AB_10596938 |
| Centrin | MPS | 04-1624 | Ms IgG2a | 1:200 | AB_10563501 |
| POC5 | TFS | PA5-24308 | Rb | 1:100 | AB_2541808 |
| SAS6 | SCBT | sc-81431 | Ms IgG2b | 1:200 | AB_1128357 |
| SAS4 | PTG | 11517-1-AP | Rb | 1:200 | AB_2244605 |
| CP110 | PTG | 12780-1-AP | Rb | 1:500 | AB_10638480 |
| CEP83 | PTG | 26013-1-AP | Rb | 1:100 | AB_2880334 |
| SCLT1 | PTG | 14875-1-AP | Rb | 1:100 | AB_2183691 |
| FBF1 | PTG | 11531-1-AP | Rb | 1:100 | AB_2246683 |
| CEP164 | SCBT | sc-515403 | Ms IgG2a | 1:100 | n/a |
| ANKRD26 | GTX | GTX128255 | Rb | 1:100 | AB_2885741 |
| TTBK2 | MPS | HPA018113 | Rb | 1:100 | AB_1858421 |
| ODF2 | MPS | HPA001874 | Rb | 1:200 | AB_1079522 |
| Centriolin | SCBT | sc-365521 | Ms IgG1 | 1:100 | AB_10851483 |
| Ninein | SCBT | sc-376420 | Ms IgG1 | 1:200 | AB_11151570 |
| CEP170 | TFS | 41-3200 | Ms IgG1 | 1:200 | AB_2533502 |
| γ-tubulin | MPS | T6557 | Ms IgG1 | 1:1000 | AB_477584 |
| γ-tubulin | MPS | T5192 | Rb | 1:1000 | AB_261690 |
| γ-tubulin | PTG | 15176-1-AP | Rb | 1:500 | AB_2878113 |
| Ac-tub | MPS | T6793 | Ms IgG2b | 1:1000 | AB_477585 |
| Arl13b | PTG | 17711-1-AP | Rb | 1:2000 | AB_2060867 |

| TREx-Confocal |  |  |  |  |  |
| --- | --- | --- | --- | --- | --- |
| SSNA1_A6 | LTK | WT#A6 | Ms IgG1 |  | n/a |
| ODF2 | Abnova | ab43840 | Rb |  | n/a |
| ATTO-NHS-594 | Atto-Tec | AD-594-31 | n/a |  | n/a |
| ExM-dSTORM |  |  |  |  |  |
| SSNA1_A6 | LTK | WT#A6 | Ms IgG1 | 1:100 | n/a |
| C2CD3 | MPS | HPA040433 | Rb | 1:200 | AB_2676981 |
| Ac-tub | TFS | 32-2700 | Ms IgG2b | 1:200 | AB_2533073 |
| Supplier Abbreviation |  |  |  |  |  |
| MPS | MilliporeSigma |  |  |  |  |
| PTG | Proteintech Group |  |  |  |  |
| TFS | Thermo Fisher Scientific |  |  |  |  |
| SCBT | Santa Cruz Biotechnology |  |  |  |  |
| R&D | R&D Systems |  |  |  |  |
| GTX | GeneTex |  |  |  |  |
| LTK | LTK BioLaboratories |  |  |  |  |
| YH | Yao-Hong Biotechnology |  |  |  |  |

## References

1. D. K. Breslow, A. J. Holland, Mechanism and Regulation of Centriole and Cilium Biogenesis. Annual Review of Biochemistry 88, 691–724 (2019).

2. J. T. Wang, T. Stearns, The ABCs of Centriole Architecture: The Form and Function of Triplet Microtubules. Cold Spring Harb Symp Quant Biol 82, 145–155 (2017).

3. G. A. Greenan, B. Keszthelyi, R. D. Vale, D. A. Agard, Insights into centriole geometry revealed by cryotomography of doublet and triplet centrioles. eLife 7, e36851 (2018).

4. S. C. Jana, G. Marteil, M. Bettencourt-Dias, Mapping molecules to structure: unveiling secrets of centriole and cilia assembly with near-atomic resolution. Current Opinion in Cell Biology 26, 96–106 (2014).

5. S. Gomes Pereira, M. A. Dias Louro, M. Bettencourt-Dias, Biophysical and Quantitative Principles of Centrosome Biogenesis and Structure. Annu. Rev. Cell Dev. Biol. 37, 43–63 (2021).

6. C. Fernandes-Mariano, J. N. Bugalhão, D. Santos, M. Bettencourt-Dias, Centrosome biogenesis and maintenance in homeostasis and disease. Current Opinion in Cell Biology 94, 102485 (2025).

7. M. LeGuennec, N. Klena, G. Aeschlimann, V. Hamel, P. Guichard, Overview of the centriole architecture. Current Opinion in Structural Biology 66, 58–65 (2021).

8. M. H. Laporte, et al., Time-series reconstruction of the molecular architecture of human centriole assembly. Cell 187, 2158–2174.e19 (2024).

9. H. Dang, E. Schiebel, Emerging roles of centrosome cohesion. Open Biology 12, 220229 (2022).

10. S. Lawo, M. Hasegan, G. D. Gupta, L. Pelletier, Subdiffraction imaging of centrosomes reveals higher-order organizational features of pericentriolar material. Nat Cell Biol 14, 1148–1158 (2012).

11. V. Mennella, et al., Subdiffraction-resolution fluorescence microscopy reveals a domain of the centrosome critical for pericentriolar material organization. Nat Cell Biol 14, 1159– 1168 (2012).

12. V. Mennella, D. A. Agard, B. Huang, L. Pelletier, Amorphous no more: subdiffraction view of the pericentriolar material architecture. Trends in Cell Biology 24, 188–197 (2014).

13. B. Madarampalli, et al., ATF5 Connects the Pericentriolar Materials to the Proximal End of the Mother Centriole. Cell 162, 580–592 (2015).

14. P. T. Conduit, A. Wainman, J. W. Raff, Centrosome function and assembly in animal cells. Nat Rev Mol Cell Biol 16, 611–624 (2015).

15. P. Gönczy, Towards a molecular architecture of centriole assembly. Nat Rev Mol Cell Biol 13, 425–435 (2012).

16. P. Guichard, V. Hamel, P. Gönczy, The Rise of the Cartwheel: Seeding the Centriole Organelle. BioEssays 40, 1700241 (2018).

17. I. Vakonakis, The centriolar cartwheel structure: symmetric, stacked, and polarized. Current Opinion in Structural Biology 66, 1–7 (2021).

18. D. Kitagawa, et al., Structural Basis of the 9-Fold Symmetry of Centrioles. Cell 144, 364– 375 (2011).

19. M. van Breugel, et al., Structures of SAS-6 Suggest Its Organization in Centrioles. Science 331, 1196–1199 (2011).

20. P. Guichard, et al., Cartwheel architecture of Trichonympha basal body. Science 337, 553 (2012).

21. Y. Lin, et al., Human microcephaly protein CEP135 binds to hSAS-6 and CPAP, and is required for centriole assembly. The EMBO Journal 32, 1141–1154 (2013).

22. D. Keller, et al., Mechanisms of HsSAS-6 assembly promoting centriole formation in human cells. Journal of Cell Biology 204, 697–712 (2014).

23. H. Gupta, et al., SAS-6 Association with γ-Tubulin Ring Complex Is Required for Centriole Duplication in Human Cells. Current Biology 30, 2395–2403.e4 (2020).

24. M. Le Guennec, et al., A helical inner scaffold provides a structural basis for centriole cohesion. Science Advances 6, eaaz4137 (2020).

25. N. Schweizer, et al., Sub-centrosomal mapping identifies augmin-γTuRC as part of a centriole-stabilizing scaffold. Nat Commun 12, 6042 (2021).

26. M. D. Arslanhan, et al., CCDC15 localizes to the centriole inner scaffold and controls centriole length and integrity. Journal of Cell Biology 222, e202305009 (2023).

27. C. Sala, et al., An interaction network of inner centriole proteins organised by POC1A-POC1B heterodimer crosslinks ensures centriolar integrity. Nat Commun 15, 9857 (2024).

28. N. A. Hall, H. Hehnly, A centriole’s subdistal appendages: contributions to cell division, ciliogenesis and differentiation. Open Biology 11, 200399 (2021).

29. J. Tischer, S. Carden, F. Gergely, Accessorizing the centrosome: new insights into centriolar appendages and satellites. Current Opinion in Structural Biology 66, 148–155 (2021).

30. D. Ma, F. Wang, J. Teng, N. Huang, J. Chen, Structure and function of distal and subdistal appendages of the mother centriole. Journal of Cell Science 136, jcs260560 (2023).

31. B. E. Tanos, et al., Centriole distal appendages promote membrane docking, leading to cilia initiation. Genes Dev. 27, 163–168 (2013).

32. G. Mazo, N. Soplop, W.-J. Wang, K. Uryu, M.-F. B. Tsou, Spatial Control of Primary Ciliogenesis by Subdistal Appendages Alters Sensation-Associated Properties of Cilia. Developmental Cell 39, 424–437 (2016).

33. N. Huang, et al., Hierarchical assembly of centriole subdistal appendages via centrosome binding proteins CCDC120 and CCDC68. Nat Commun 8, 15057 (2017).

34. T. T. Yang, et al., Super-resolution architecture of mammalian centriole distal appendages reveals distinct blade and matrix functional components. Nat Commun 9, 2023 (2018).

35. W. M. Chong, et al., Super-resolution microscopy reveals coupling between mammalian centriole subdistal appendages and distal appendages. eLife 9, e53580 (2020).

36. T.-J. B. Chang, J. C.-C. Hsu, T. T. Yang, Single-molecule localization microscopy reveals the ultrastructural constitution of distal appendages in expanded mammalian centrioles. Nat Commun 14, 1688 (2023).

37. D. Kumar, J. Reiter, How the centriole builds its cilium: of mothers, daughters, and the acquisition of appendages. Current Opinion in Structural Biology 66, 41–48 (2021).

38. S. Shakya, C. J. Westlake, Recent advances in understanding assembly of the primary cilium membrane. Fac Rev 10, 16 (2021).

39. H. Zhao, Z. Khan, C. J. Westlake, Ciliogenesis membrane dynamics and organization. Seminars in Cell & Developmental Biology 133, 20–31 (2023).

40. X. Ye, H. Zeng, G. Ning, J. F. Reiter, A. Liu, C2cd3 is critical for centriolar distal appendage assembly and ciliary vesicle docking in mammals. Proceedings of the National Academy of Sciences 111, 2164–2169 (2014).

41. J.-J. Tsai, W.-B. Hsu, J.-H. Liu, C.-W. Chang, T. K. Tang, CEP120 interacts with C2CD3 and Talpid3 and is required for centriole appendage assembly and ciliogenesis. Sci Rep 9, 6037 (2019).

42. A. Nommick, et al., Lrrcc1 and Ccdc61 are conserved effectors of multiciliated cell function. Journal of Cell Science 135, jcs258960 (2022).

43. N. Gaudin, et al., Evolutionary conservation of centriole rotational asymmetry in the human centrosome. eLife 11, e72382 (2022).

44. M. H. Laporte, et al., Human SFI1 and Centrin form a complex critical for centriole architecture and ciliogenesis. The EMBO Journal 41, e112107 (2022).

45. F. Ramos-Morales, C. Infante, C. Fedriani, M. Bornens, R. M. Rios, NA14 Is a Novel Nuclear Autoantigen with a Coiled-coil Domain*. Journal of Biological Chemistry 273, 1634–1639 (1998).

46. F. Pfannenschmid, et al., Chlamydomonas DIP13 and human NA14: a new class of proteins associated with microtubule structures is involved in cell division. Journal of Cell Science 116, 1449–1462 (2003).

47. J. Schoppmeier, W. Mages, K.-F. Lechtreck, GFP as a tool for the analysis of proteins in the flagellar basal apparatus of Chlamydomonas. Cell Motility 61, 189–200 (2005).

48. H. P. Price, et al., The Orthologue of Sjögren’s Syndrome Nuclear Autoantigen 1 (SSNA1) in Trypanosoma brucei Is an Immunogenic Self-Assembling Molecule. PLoS ONE 7, e31842 (2012).

49. M. F. Lévêque, L. Berry, S. Besteiro, An evolutionarily conserved SSNA1/DIP13 homologue is a component of both basal and apical complexes of *Toxoplasma gondii*. Scientific Reports 6, 27809 (2016).

50. A. Errico, P. Claudiani, M. D’Addio, E. I. Rugarli, Spastin interacts with the centrosomal protein NA14, and is enriched in the spindle pole, the midbody and the distal axon. Human Molecular Genetics 13, 2121–2132 (2004).

51. U. Goyal, B. Renvoisé, J. Chang, C. Blackstone, Spastin-Interacting Protein NA14/SSNA1 Functions in Cytokinesis and Axon Development. PLoS One 9 (2014).

52. M. Rodríguez-Rodríguez, et al., Characterization of the structure and self-recognition of the human centrosomal protein NA14: implications for stability and function. *Protein Engineering*, Design and Selection 24, 883–892 (2011).

53. N. Basnet, et al., Direct induction of microtubule branching by microtubule nucleation factor SSNA1. Nat Cell Biol 20, 1172–1180 (2018).

54. 54. L. Agostini, et al., Structural insights into SSNA1 self-assembly and its microtubule binding for centriole maintenance. [Preprint] (2024). Available at: https://www.biorxiv.org/content/10.1101/2024.11.13.623454v1 [Accessed 25 April 2025].

55. E. J. Lawrence, G. Arpag, C. Arnaiz, M. Zanic, SSNA1 stabilizes dynamic microtubules and detects microtubule damage. eLife 10, e67282 (2021).

56. T. Aki, T. Funakoshi, J. Nishida-Kitayama, Y. Mizukami, TPRA40/GPR175 regulates early mouse embryogenesis through functional membrane transport by Sjögren’s syndrome-associated protein NA14. Journal of Cellular Physiology 217, 194–206 (2008).

57. G. D. Amatngalim, et al., Measuring cystic fibrosis drug responses in organoids derived from 2D differentiated nasal epithelia. Life Science Alliance 5 (2022).

58. H. G. Damstra, et al., Visualizing cellular and tissue ultrastructure using Ten-fold Robust Expansion Microscopy (TREx). eLife 11, e73775 (2022).

59. 59. W. Nijenhuis, et al., Optical nanoscopy reveals SARS-CoV-2-induced remodeling of human airway cells. [Preprint] (2021). Available at: https://www.biorxiv.org/content/10.1101/2021.08.05.455126v1 [Accessed 5 January 2025].

60. R. Rashpa, M. Brochet, Expansion microscopy of Plasmodium gametocytes reveals the molecular architecture of a bipartite microtubule organisation centre coordinating mitosis with axoneme assembly. PLOS Pathogens 18, e1010223 (2022).

61. S. Li, et al., ELI trifocal microscope: a precise system to prepare target cryo-lamellae for in situ cryo-ET study. Nat Methods 20, 276–283 (2023).

62. T. Oda, H. Yanagisawa, R. Kamiya, M. Kikkawa, A molecular ruler determines the repeat length in eukaryotic cilia and flagella. Science 346, 857–860 (2014).

63. M. Ma, et al., Structure of the Decorated Ciliary Doublet Microtubule. Cell 179, 909–922.e12 (2019).

64. J.-H. Wei, Z. C. Zhang, R. M. Wynn, J. Seemann, GM130 Regulates Golgi-Derived Spindle Assembly by Activating TPX2 and Capturing Microtubules. Cell 162, 287–299 (2015).

65. M. Mirdita, et al., ColabFold: making protein folding accessible to all. Nat Methods 19, 679– 682 (2022).

66. L. Wang, M. Failler, W. Fu, B. D. Dynlacht, A distal centriolar protein network controls organelle maturation and asymmetry. Nat Commun 9, 3938 (2018).

67. E. M. Park, et al., WBP11 is required for splicing the TUBGCP6 pre-mRNA to promote centriole duplication. Journal of Cell Biology 219, e201904203 (2019).

68. C. Thauvin-Robinet, et al., The oral-facial-digital syndrome gene C2CD3 encodes a positive regulator of centriole elongation. Nat Genet 46, 905–911 (2014).

69. A. N. Hoover, et al., C2cd3 is required for cilia formation and Hedgehog signaling in mouse. Development 135, 4049–4058 (2008).

70. C. R. Cortés, et al., Mutations in human C2CD3 cause skeletal dysplasia and provide new insights into phenotypic and cellular consequences of altered C2CD3 function. Sci Rep 6, 24083 (2016).

71. 71. J. A. Pfister, et al., The C. elegans homolog of Sjögren’s Syndrome Nuclear Antigen 1 is required for the structural integrity of the centriole and bipolar mitotic spindle assembly. [Preprint] (2024). Available at: https://www.biorxiv.org/content/10.1101/2024.10.03.616528v1 [Accessed 25 April 2025].

72. Z. C. Zhu, et al., Interactions between EB1 and Microtubules. Journal of Biological Chemistry 284, 32651–32661 (2009).

73. H. Inaba, et al., Generation of stable microtubule superstructures by binding of peptide-fused tetrameric proteins to inside and outside. Science Advances 8, eabq3817 (2022).

74. L. von Tobel, et al., SAS-1 Is a C2 Domain Protein Critical for Centriole Integrity in C. elegans. PLOS Genetics 10, e1004777 (2014).

75. A. Woglar, et al., Molecular architecture of the C. elegans centriole. PLOS Biology 20, e3001784 (2022).

76. 76. P. Gönczy, et al., C. elegans SAS-1 ensures centriole integrity and ciliary function, and operates with SSNA-1. [Preprint] (2025). Available at: https://www.biorxiv.org/content/10.1101/2025.04.22.650004v1 [Accessed 25 April 2025].

77. I. A. Vorobjev, Yu. S. Chentsov, The ultrastructure of centriole in mammalian tissue culture cells. Cell Biology International Reports 4, 1037–1044 (1980).

78. C. K. Lai, et al., Functional characterization of putative cilia genes by high-content analysis. MBoC 22, 1104–1119 (2011).

79. S. Xie, N. Naslavsky, S. Caplan, Emerging insights into CP110 removal during early steps of ciliogenesis. Journal of Cell Science 137, jcs261579 (2024).

80. J. F. Reiter, M. R. Leroux, Genes and molecular pathways underpinning ciliopathies. Nat Rev Mol Cell Biol 18, 533–547 (2017).

81. A. Micsonai, et al., Accurate secondary structure prediction and fold recognition for circular dichroism spectroscopy. Proceedings of the National Academy of Sciences 112, E3095– E3103 (2015).

